# Abundantly expressed class of non-coding RNAs conserved through the multicellular evolution of dictyostelid social amoebae

**DOI:** 10.1101/2020.04.05.026054

**Authors:** Jonas Kjellin, Lotta Avesson, Johan Reimegård, Zhen Liao, Ludwig Eichinger, Angelika Noegel, Gernot Glöckner, Pauline Schaap, Fredrik Söderbom

## Abstract

**Background:** Aggregative multicellularity has evolved multiple times in diverse groups of eukaryotes. One of the most well-studied examples is the development of dictyostelid social amoebae, e.g. *Dictyostelium discoideum*. However, it is still poorly understood why multicellularity emerged in these amoebae while the great majority of other members of Amoebozoa are unicellular. Previously a novel type of non-coding RNA, Class I RNAs, was identified in *D. discoideum* and demonstrated to be important for normal multicellular development. In this study we investigated Class I RNA evolution and its connection to multicellular development.

**Results:** New Class I RNA genes were identified by constructing a co-variance model combined with a scoring system based on conserved up-stream sequences. Multiple genes were predicted in representatives of each major group of Dictyostelia and expression analysis validated that our search approach can identify expressed Class I RNA genes with high accuracy and sensitivity. Further studies showed that Class I RNAs are ubiquitous in Dictyostelia and share several highly conserved structure and sequence motifs. Class I RNA genes appear to be unique to dictyostelid social amoebae since they could not be identified in searches in outgroup genomes, including the closest known relatives to Dictyostelia.

**Conclusion:** Our results show that Class I RNA is an ancient abundant class of ncRNAs, likely to have been present in the last common ancestor of Dictyostelia dating back at least 600 million years. Taken together, our current knowledge of Class I RNAs suggests that they may have been involved in evolution of multicellularity in Dictyostelia.

## Background

The role of RNA goes far beyond it being an intermediate transmitter of information between DNA and protein, in the form of messenger (m)RNAs. This has been appreciated for a long time for some non-coding RNAs (ncRNAs), such as transfer (t)RNAs, ribosomal (r)RNAs, small nuclear (sn)RNAs, and small nucleolar (sno)RNAs. Today, we know that ncRNAs are involved in regulating most cellular processes and the advent of high-throughput sequencing technologies have facilitated the identification of numerous different classes of ncRNAs [1]. These regulatory RNAs vary greatly in size from 21-24 nucleotides (nt), e.g. micro (mi)RNAs and small interfering (si)RNAs, to several thousands of nucleotides, such as long non-coding (lnc)RNAs. Several classes of ncRNAs are ubiquitously present in all domains of life while others are specific to certain evolutionary linages, contributing to their specific characteristics. This can be exemplified by ncRNAs in Metazoa, where an increase in number of ncRNAs, e.g. miRNAs, is associated with increased organismal complexity and is believed to be essential for the evolution of metazoan multicellularity [2]. Multicellularity in plants and animals is achieved by clonal division and development originating from a single cell. This is in contrast to aggregative multicellularity, were cells stream together to form multicellular structures upon specific environmental changes. Aggregative multicellularity has evolved independently multiple times and is found both among eukaryotes and prokaryotes [3–10]. The complexity of the aggregative multicellular life stages varies for different organisms, but they all share the transition from unicellularity to coordinated development upon environmental stress, e.g. starvation, which eventually leads to formation of fruiting bodies containing cysts or spores [3]. Probably the most well-studied aggregative multicellularity is the development of the social amoeba *Dictyostelium discoideum* belonging to the group Dictyostelia within the supergroup Amoebozoa. Dictyostelia is a monophyletic group estimated to date back at least 600 million years [11], which is similar to the age of Metazoa [12]. Dictyostelia is currently divided into four major groups (Group 1-4) where all members share the ability of transition from uni-to multicellularity upon starvation [13]. However, the complexity of the development and the morphology of the fruiting bodies varies between different dictyostelids, where the highest level of multicellular complexity is found among group 4 species, which includes *D. discoideum* [14, 15]. Recently a new taxonomy was proposed for many dictyostelids [16]. As this new taxonomy has not yet been fully adopted by the research community, we choose to use the previous designations throughout this study (old and new names, including NCBI accession numbers, are summarized in Additional file 1).

Well-annotated genome sequences are available for representative species of all four major groups of Dictyostelia [11, 17–20] and multiple draft genome sequences are available for additional dictyostelids. This has allowed for comparative genomics, which has provided information about the protein coding genes that are important for the diversification of Dictyostelia from other amoebozoans. Comparison between genomes has also given insight into the genes required for the evolution of the distinct morphological characteristics, which define each group [11, 17, 18, 20, 21]. However, evolution of complex traits such as multicellularity in Dictyostelia as well as other eukaryotic groups, cannot solely be explained by the appearance of novel genes but also relies on an increased ability to regulate pre-existing genes and their products so that they can function in novel genetic networks [20, 22]. This is also supported by the major transcriptional reprogramming during multicellular development in *D. discoideum* [23].

*D. discoideum* harbors several classes of developmentally regulated ncRNAs with regulatory potential, e.g. microRNAs [24–27], long non-coding RNAs [28] and long antisense RNAs [28, 29]. In addition, a large part of the ncRNA repertoire of *D. discoideum* is constituted by Class I RNAs, originally identified in full length cDNA libraries [30]. So far, Class I RNAs have only been validated in *D. discoideum* [30, 31], but they have also been computationally predicted in *Dictyostelium purpureum* [18]. Both species belong to the same evolutionary group of Dictyostelia, i.e. group 4 [13, 16]. In *D. discoideum*, Class I RNAs are 42-65 nt long and are expressed at high levels from a large number of genes. Members of Class I RNAs are characterized by a short stem structure, connecting the 5’ and 3’ ends, and a conserved 11 nt sequence motif adjacent to the 5’part of the stem. The remainder of the RNA is variable both in sequence and structure. Class I RNAs mainly localize to the cytoplasm [30] where one of the RNAs has been shown to associate with four different proteins of which at least one, the RNA recognition motifs (RRM) containing protein CIBP, directly binds to the Class I RNA [31]. Furthermore, the Class I RNAs appear to be involved in regulating multicellular development. This is based on the observations that Class I RNAs are developmentally regulated and that cells where a single Class I RNA gene has been knocked out show aberrant early development [30, 31].

In this study we investigated the prevalence of Class I RNAs within Dictyostelia as well as in other organisms with the aim to understand if Class I RNAs are restricted to dictyostelids and perhaps required for their aggregative multicellularity. Based on the known Class I RNAs from *D. discoideum*, we know that the major part of the RNA is variable and hence sequence-based searches alone, such as BLAST, would not reliably identify new genes. This was solved by constructing a Class I classifier based on a co-variance model, which includes both sequence and structure information, combined with a scoring system for conserved up-stream sequence motifs, e.g. promoter motifs. Using this search approach, we identified approximately 300 Class I RNA genes predicted to be expressed in the genomes of 16 different dictyostelids. In addition, we validate the expression of ~100 Class I RNAs from six different species, using both northern blot and RNA-seq, which supports the high accuracy and sensitivity of our search approach to identify expressed Class I RNAs. Comparative studies of identified Class I RNA loci show several well conserved features, like stem forming properties connecting the 5’ and 3’ ends, as well as preserved sequence motifs. Importantly, Class I RNAs appear to be specific to Dictyostelia as no Class I RNA genes were identified in genomes of organisms outside this group of social amoebae, including their closest known relatives or organisms exhibiting different kinds of multicellularity. Taken together, this unique class of ncRNAs constitute a very large number of conserved highly expressed genes involved in the evolution of Dictyostelia multicellular development.

## Results

### Co-variance model identifies Class I RNA genes in evolutionary distinct groups of Dictyostelia social amoebae

We have previously identified and characterized Class I RNAs from the social amoeba *D. discoideum*. Members of Class I RNAs are characterized by a short stem-structure connecting the 5’ and 3’ ends and a conserved 11 nt sequence motif adjacent to the 5’part of the stem. The remainder of the Class I RNAs are highly variable both in sequence and structure (Fig. 1a). This class of ncRNA have so far only been validated in *D. discoideum* where it is associated with development [30, 31], but has also been predicted in *D. purpureum* [18].

**Figure 1.**
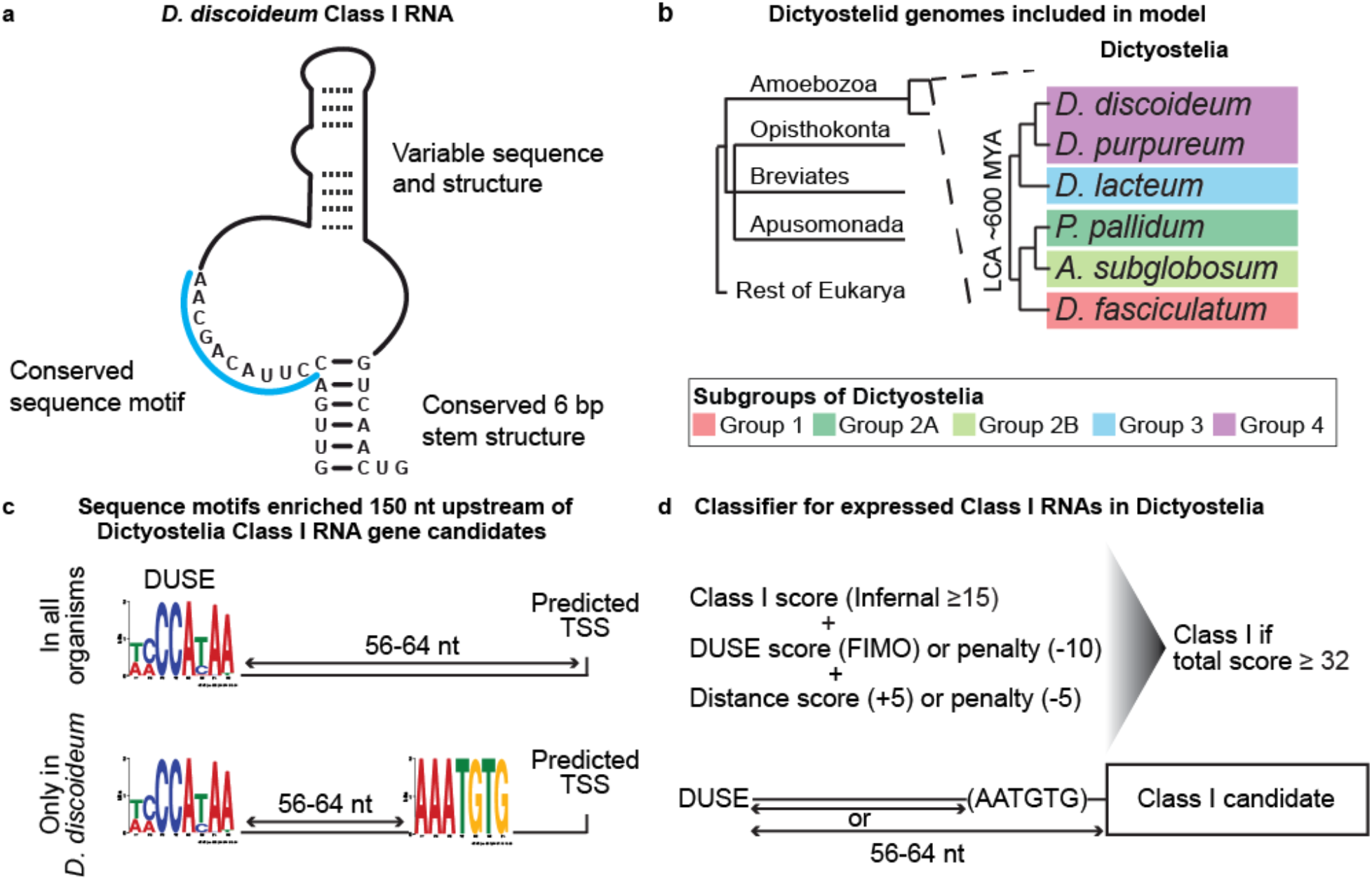
Search strategy and classification of Class I RNAs. a) Schematic representation of previously described *D. discoideum* Class I RNAs [31]. b) Schematic phylogeny showing the location of Amoebozoa, a sister group to Obazoa (Opisthokonta, Breviates and Apusomonada) in the eukaryotic tree of life based on [35]. Dictyostelia is represented by species belonging to each major group [13]. The genomes of these dictyostelids were searched for Class I RNA genes and newly identified genes were used to refine the co-variance model. c) Enriched sequence motifs identified upstream of Class I RNA gene candidates in the different dictyostelids represented in panel b (Infernal score ≥ 25, n=126). The putative promoter motif (DUSE) is found approximately 60 nt from the predicted start of transcription (TSS) in all organisms (upper part). DUSE in combination with TGTG-box, only identified in *D. discoideum* (lower part). d) Summary of scoring system used for the classifier of Dictyostelia Class I RNA based on Infernal score ≥ 15, presence of DUSE, and distance between DUSE and predicted TSS or TGTG-box.

The presence of Class I RNA genes in two different dictyostelids and the fact that at least one Class I RNA member is involved in controlling early multicellular development, led us to hypothesize that this class of ncRNA may be a general effector for early development in all members of dictyostelid social amoebae. Hence, it may have been present in the last common ancestor of Dictyostelia, dating back approximately 600 million years in evolution. In order to investigate this, we used the complete and well-annotated genome sequences for representatives of each major group of Dictyostelia, i.e. *D. discoideum* [17], *D. purpureum* [18], *Dictyostelium lacteum* [20], *Polysphondylium pallidum* [11], *Acytostelium subglobosum* [19] and *Dictyostelium fasciculatum* [11] (Fig. 1b). Class I RNAs cannot be reliably detected by sequence searches alone due to the high sequence variability. Therefore, we constructed a co-variance model (CM) with Infernal [32] where both sequence and secondary structure information of 34 *D. discoideum* Class I RNAs were taken into account (see Materials and methods for details). The initial CM search of the six Dictyostelia genomes, followed by manual inspection of the results, indicated the presence of Class I RNAs in all major groups of Dictyostelia (Additional file 2: Fig. S1). These candidates scored ≥ 25 in the CM search and had the potential to form a short stem similar to the *D. discoideum* Class I RNAs. In order to improve the CM, these candidates (CM score ≥ 25 and potential to form stem) were added to the CM followed by new genome searches. This process was repeated until no new candidates fulfilling the criteria were identified after which a final search with increased sensitivity was performed (see Materials and methods). In total, 126 loci distributed over all major groups of Dictyostelia were identified including 36 of the 40 published *D. discoideum* Class I RNAs [30, 31] and all the 26 previously predicted *D. purpureum* Class I loci [18] (Additional file 2: Fig. S1).

### Refining the search for Class I RNA genes using conserved promoter elements

Many *D. discoideum* ncRNA genes have an upstream putative promoter element, DUSE (*Dictyostelium* upstream sequence element), which in most cases is situated ~60 nt from the transcriptional start site (TSS) [33]. However, for *D. discoideum* Class I RNAs, DUSE is often found further upstream. In these cases, a TGTG-box (AAATGTG) is located ~60 nt downstream of DUSE while the distance from the start of the mature RNA varies. Whether the TGTG-box is an additional promotor element or the TSS of a precursor transcript is currently not known. DUSE appears to be conserved within group 4 of Dictyostelia, as it was also identified ~60 nt in front of the predicted *D. purpureum* Class I RNAs [18]. In order to investigate the presence of conserved upstream motifs in the rest of Dictyostelia, we searched for enriched motifs in the 150 nt upstream sequence of all the 126 Class I RNA gene candidates identified in the CM search. Intriguingly, DUSE like motifs could be identified ~60 nt upstream of the predicted start for the majority of the Class I RNA gene candidates (73 of 126) in all organisms. In contrast, the TGTG-box was only found in a subset (21 of 126) of the upstream sequences of which all belonged to *D. discoideum* Class I RNA genes (Fig. 1c). As both sequence and distance of DUSE appeared to be conserved in all major groups of Dictyostelia, we used this information to create a scoring system anticipating accurate prediction of expressed Class I RNA genes. (Fig. 1d). First the score produced by the CM search (Infernal) was used and all candidates with a score ≥ 15 were included in order to capture more divergent Class I RNA genes. Next we scored the presence and location of DUSE and TGTG-box 150 nt upstream of the candidates identified in the CM search based on the motif identification program FIMO [34]. Lack of DUSE and/or non-canonical distance from predicted TSS or TGTG-box was penalized with negative scores. Taken together, a total score of 32 could be achieved if a high-scoring DUSE was identified at the predicted distance upstream of a candidate Class I RNA gene with the lowest allowed Infernal score (≥ 15). Based on this, all candidates scoring 32 or higher were classified as Class I RNA loci. Using this approach, we predicted 18-39 Class I RNAs (146 in total) for each of the six dictyostelids investigated (see below).

### Class I RNAs of predicted sizes are expressed at high levels in all four groups of Dictyostelia

Based on the Class I classifier, we predicted Class I RNA genes in all dictyostelids included in the CM build. But are all these genes really expressed and how accurate are the size predictions? From our previous studies, we know that Class I RNAs in *D. discoideum* are expressed at high levels at vegetative growth and are readily detected by northern blot [30, 31]. Hence, we used the same approach to validate a subset of randomly chosen candidates that made the total score threshold in *D. purpureum* (Group 4), *D. lacteum* (Group 3), *P. pallidum* (Group 2A), *A. subglobosum* (Group 2B) and *D. fasciculatum* (Group 1). RNA was prepared from vegetative growing amoebae and specific Class I RNAs were analyzed by northern blot (Fig. 2a). For *D. purpureum*, we probed for DpuR-7, predicted to be 54 nt long, resulting in a strong signal. In addition, we designed two probes predicted to recognize six different 85 nt long RNAs (DpuR-X) and the majority (24/30) of Class I RNAs (DpuR-Y), respectively. As expected, probing for DpuR-X resulted in one band on the northern blot. For DpuR-Y, we expected signals from several Class I RNAs to overlap due to similar/identical sizes but we could still detect several distinct bands within the expected size range. In *D. fasciculatum*, we probed for one Class I RNA predicted to be 62 nt long (DfaR-4) while at least two candidates were probed for in the other organisms, i.e. *D: lacteum:* DlaR-1 (61 nt) and DlaR-5 (54 nt), *P. pallidum:* PpaR-1/2 (59/62 nt) and PpaR-9 (58 nt), and *A. subglobosum:* AsuR-13/14 (61 nt) and AsuR-10 (82 nt). Distinct bands were detected for each Class I RNA candidate and the sizes matched the predictions well although the northern results often indicated that the RNAs were a few nt longer than predicted (Fig. 2a and see below). The larger (but much weaker) signals observed for AsuR-13/14 and PpaR-9 are likely cross hybridizations to longer Class I RNAs. Taken together, the results confirm that Class I RNAs are conserved and expressed in all four groups of Dictyostelia social amoebae.

**Figure 2.**
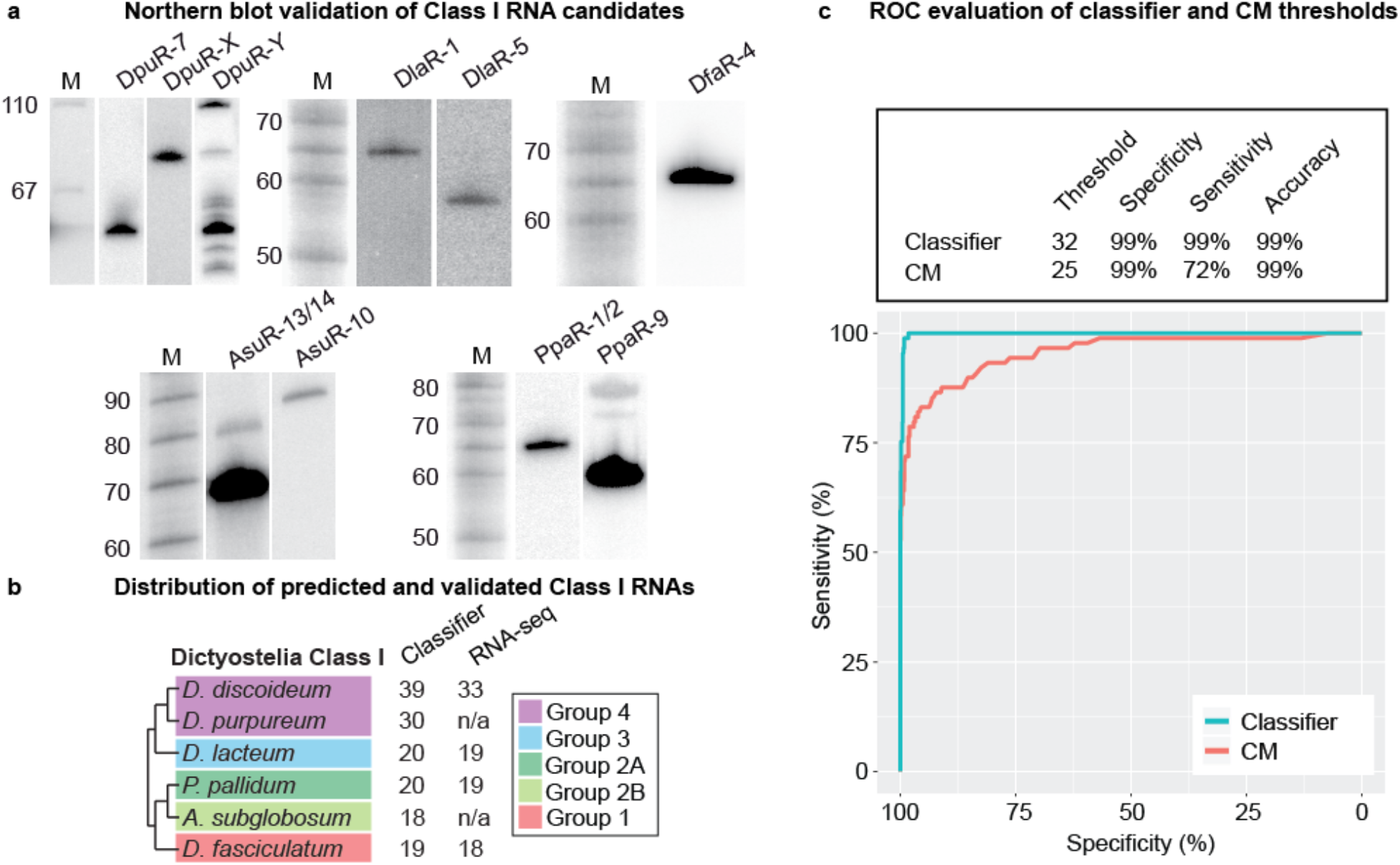
Expression of predicted Class I RNA genes. a) Northern blot validation of different Class I RNAs from *D. purpureum* (DpuR), *D. lacteum* (DlaR), *D. fasciculatum* (DfaR), *A. subglobosum* (AsuR), and *P. pallidum* (DpaR). The number after the species-specific designations indicates which Class I RNA the probe recognizes. When two numbers are given, the probe recognize two different Class I RNAs. DpuR-X indicates that the probe is expected to hybridize to six different Class I RNAs predicted to be 85 nt long. DpuR-Y indicates that the probe is expected to hybridize to 24 Class I RNAs. For each organism except for *D. fasciculatum* (DfaR), the same membrane was probed, stripped, and reprobed for the different Class I RNAs. Radioactively labeled size marker is indicated by M and numbers to the left indicate sizes in nucleotides. b) Number of Class I RNA genes in each species according to the classifier. RNA-seq designate the number of expressed Class I RNA genes verified by RNA-seq. c) ROC curves based on the RNA-seq validation and either classifier score or CM score for all Class I candidates in *D. discoideum, D. lacteum, P. pallidum*, and *D. fasciculatum* identified in the CM search. Input data available in Additional file 3. Evaluation of the classifier and CM thresholds used throughout the study are shown above the plot. Individual ROC curves for each organism are found in Additional file 2: Fig. S2.

### Classifier accurately predicts expressed Class I RNAs in all major groups of Dictyostelia

Next, we performed RNA-seq on *D. discoideum, D. lacteum, P. pallidum* and *D. fasciculatum* representing each major group of Dictyostelia. RNA was prepared from growing cells as well as two multicellular life-stages, i.e. mound and slug/finger stages, to increase our chances to also detect Class I RNAs that are only expressed at specific life stages. Expression was evaluated based on the read count and coverage over all loci identified in the CM search (Infernal score ≥ 15) as exemplified in Additional file 2: Fig. S2. Strikingly, expression both during vegetative growth and development could be confirmed for almost all Class I RNA candidates identified by the classifier (total score ≥ 32) (Fig. 2b, Additional file 3). Next, we calculated receiver operator characteristics (ROC) curves in order to evaluate the classifier performance and investigate if it improves Class I RNA identification compared to CM search alone. ROC curves were generated for *D. discoideum, D. lacteum, P. pallidum* and *D. fasciculatum* individually (Additional file 2: Fig. S3) as well as for the pooled data (Fig. 2c) based on the RNA-seq validation and either CM search score (≥ 25) or classifier score (≥ 32). Evaluation of the two search approaches show an increase in both sensitivity and accuracy of prediction for the classifier, i.e. when the promoter (DUSE) presence and distance were included in the classification of Class I RNA gene candidates. To summarize, the classifier reliably detects expressed Class I RNAs in all the tested dictyostelids with almost no false positives.

### Conserved features of Dictyostelia Class I RNA

The wealth of newly identified Class I RNA genes in six different and evolutionary separated dictyostelids allowed us to construct a general/unifying picture of Class I RNAs. This will also be of importance when searching for Class I RNA genes in other species in order to track down the birth of this class of ncRNAs in evolution (see below).

#### Class I transcription is dependent on DUSE

Both the sequence of DUSE and its upstream location is highly conserved in all of the analyzed amoebae (Fig. 3a). In the group 1 and 4 dictyostelids, DUSE contains three consecutive C residues, while two consecutive C’s are found in the majority of the DUSE of *A. subglobosum* (Group 2B) and *D. lacteum* (Group 3) and in all *P. pallidum* (Group 2A). The RNA-seq data show that the promoter element is essential for transcription as it is found in front of all expressed Class I RNA genes. Further strengthening its importance is the observation that high scoring Class I RNAs (both considering score produced by classifier or CM alone) lacking DUSE at the correct up-stream location are not expressed (Additional file 3). The TGTG-box, situated 60 nt down-stream of the DUSE and in front of the predicted TSS, is only found in *D. discoideum* suggesting that this is a rather late addition in the evolution of Class I RNA genes. The genes are most likely transcribed by RNA polymerase III (Pol III) as the majority of the Class I RNAs from the different dictyostelids exhibit a stretch of at least four consecutive T-residues downstream of the predicted end of transcription, which is a common Pol III termination signal [36].

**Figure 3.**
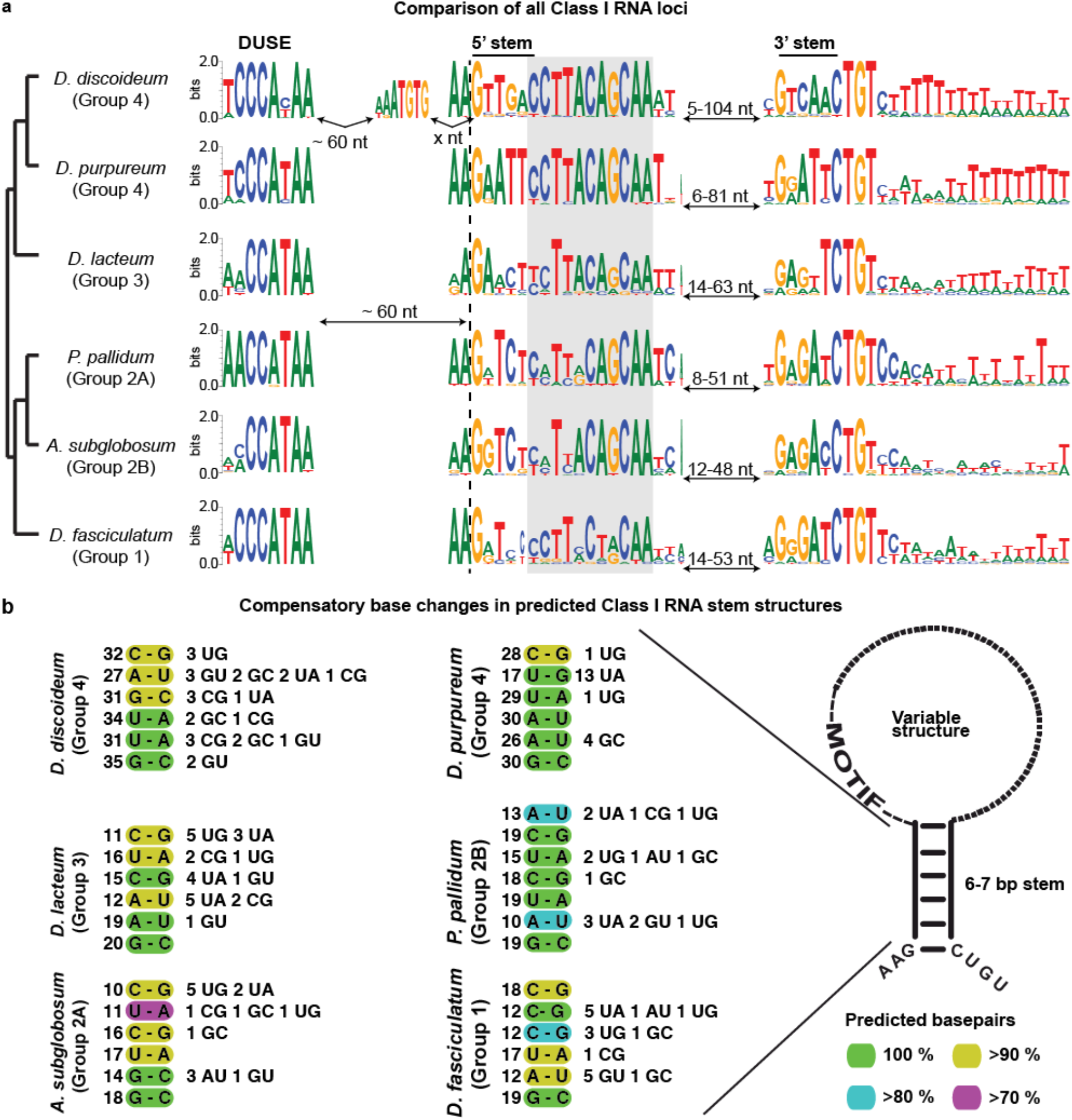
Conserved characteristics of Class I RNAs. a) Sequence logo representing species specific alignments of conserved features of Class I RNA loci. DUSE indicate the putative promoter element and the 5’ and 3’stem sequences of the conserved stem-structure are indicated. The conserved 11 nt sequence motif adjacent to the 5’part of the stem is boxed in gray. The dashed vertical line denotes the predicted start of the Class I RNAs based on the CM search. The sequence logo between DUSE and the 5’ stem motif for *D. discoideum* represent the TGTG-box. Numbers of nucleotides (nt) correspond to the distances between indicated motifs. b) Displayed are nucleotides representing the most common base pair for each position (numbers to the left) of the conserved stem structure for each organism (color key down right). Numbers to the right represent less common nucleotide combinations predicted to base pair. The few combinations of nucleotides not predicted to base pair are not shown. Schematic structure of Class I RNA is presented to the right.

#### 5’ and 3’ ends of Class I RNAs are conserved

The RNA-seq analyses showed that the *D. discoideum* Class I RNAs started at the predicted (and previously defined [30]) G residue while the majority of group 1-3 Class I RNAs started one to two nucleotides upstream of the conserved G (Additional file 2: Fig. S2). When we compared all loci, we noticed that the two nucleotides preceding the completely conserved G residue, are highly conserved A-residues in all six species investigated. The difference in 5’ ends of the mature Class I RNAs indicates that either transcription initiation or 5’ processing of Class I RNAs differ between group 4 species and species belonging to the other evolutionary groups of Dictyostelia. Also, the very 3’ end of Class I RNAs is highly similar with an almost perfectly conserved CTGT sequence in the genomic loci. The coverage from the RNA-seq data indicates that these four nucleotides are transcribed so that the CUGU sequence is included in the mature Class I RNA (Additional file 2: Fig. S2) where the C residue always have base pairing potential with the conserved 5’ G residue (Fig. 3a-b). The RNA-seq coverage agrees with the slightly longer than predicted lengths of Class I RNAs observed by northern blot (see above).

#### Class I RNA GC-content, size, sequence motif, and stem-structure are conserved throughout Dictyostelia

Comparison of all identified Class I RNAs showed that both Class I RNA length (median of ~60 nt) and GC content (32-41%) are highly conserved (Additional file 2: Fig. S4a-b). The stable GC content of Class I RNAs is remarkable considering the variation in overall genome GC content for these six dictyostelids (Additional file 2: Fig. S4b). In spite of these conserved features and specific sequence motifs discussed above and later, the overall sequence variability of Class I RNAs is extensive both within and between species. Only a few examples of loci with identical sequences are found within *D. discoideum, D. purpureum* and *P. pallidum*, respectively (Additional file 3). No Class I genes with identical sequences were found between these six analyzed species.

In contrast to the overall variable sequences, the 11 nt sequence motif identified among the *D. discoideum* Class I RNAs, is highly conserved both within and between all six dictyostelids (marked in grey in Fig. 3a). The motif is nearly perfectly conserved within group 4, while some positions of the motif are variable in group 1-3. However, T, C, C, A, and A at position 3, 6, 9, 10 and 11 (counting from the 5’most nt of the motif) are almost identical between all the Class I RNAs regardless of species. Other nucleotides are well conserved in most of the evolutionary groups but not all. The sequence motif does not seem to extensively engage in base-pairing since computational prediction indicates that the conserved motif is less structured compared to the full-length RNA in most of the organisms (Additional file 2: Fig. S4c-d). However, the first 5’-nucleotide of the motif is often part of the stem-structure (see below), while the base pairing potential for the remainder of the sequence drops in a pattern similar for all six dictyostelids (Additional file 2: Fig. S4e).

Another distinct feature common to all *D. discoideum* Class I RNAs is the short (six bp) stem structure, connecting the 5’ and 3’ ends of the RNA (Fig. 3a). This stem is predicted to be present in all Class I RNAs in all six species. However, in contrast to the conserved sequence motif, the nucleotide sequence of the stem-structure has changed substantially during Dictyostelia evolution. Nevertheless, the base pairing potential is retained, indicating that it is the structure rather than sequence that is crucial for function (Fig. 3b). This is further supported by the high number of compensatory mutations found in the predicted stem of Class I RNAs within each species (Fig. 3b). Notably, in spite of the sequence variation in the stems, the 5’ most G is completely conserved within all Class I RNAs from all six dictyostelids representing each evolutionary group of Dictyostelia. The predicted base-paired structure of the stem and the unstructured feature of the conserved sequence motif correspond well with previous *in vitro* probing results of one Class I RNA, DdR21, from *D. discoideum* [31]. Taken together, Class I GC-content, length, stem structure and 11 nt motif are highly conserved in all evolutionary groups of Dictyostelia, indicating that these parts are essential for Class I RNA function.

### Class I RNAs are developmentally regulated and highly conserved throughout Dictyostelia

We knew from our previous work that *D. discoideum* Class I RNAs are developmentally regulated and that Class I knock-out cells, *DdR-21* k.o., are disturbed in early development, leading to more and smaller fruiting bodies compared to wild-type cells [31]. In order to investigate if the developmental regulation is conserved also in other dictyostelids, we performed principal component analysis (PCA) of Class I expression in *D. discoideum, P. pallidum* and *D. fasciculatum* based on the RNA-seq data. *D. lacteum* was not included in this analysis since only one replicate per timepoint was available. The PCA plots show developmental regulation of Class I RNAs in all three amoebae as the different life stages are clearly separated (Additional file 2: Fig. S5). Taken together, this suggests a role for Class I RNAs in regulating multicellular development in Dictyostelia.

If the prediction that Class I RNAs are involved in and important for multicellular development holds true, these ncRNAs should be present in all dictyostelids. To analyze this, we used the Class I classifier to investigate the presence of Class I RNA genes in ten additional social amoebae genome sequences. Class I RNA genes were detected in all species, 9-31 genes in each genome, of which the great majority passed manual curation based on the ability to form a short stem connecting the 5’ and 3’ end (Fig. 4). It should be noted that these are draft genome sequences of varying degree of completeness (Additional file 1). Comparison of all curated loci reinforced the previously identified conserved Class I features i.e. the high sequence conservation of the terminal residues and the 11 nt motif as well as the short stem where structure but not sequence is preserved. In addition, presence of DUSE ~60 nt up-stream of the majority of the identified Class I RNA genes strongly suggests that expressed Class I RNAs exist in all members of Dictyostelia. The TGTG-box, previously only found upstream of *D. discoideum* Class I RNA loci, was identified in four additional genomes all belonging to group 4 or the *P. violaceum* complex strengthening the hypothesis that this motif emerged rather late in Class I evolution. In addition, both Class I RNA lengths and GC content are conserved also when considering all species (Additional file 2: Fig. S6). Taken together, this proves the existence and emphasize the importance of Class I RNA genes throughout the evolution of Dictyostelia social amoebae. We have named all curated Class I RNA genes according to the naming convention previously defined for *D. discoideum* Class I RNA genes [30] (Additional file 4).

**Figure 4.**
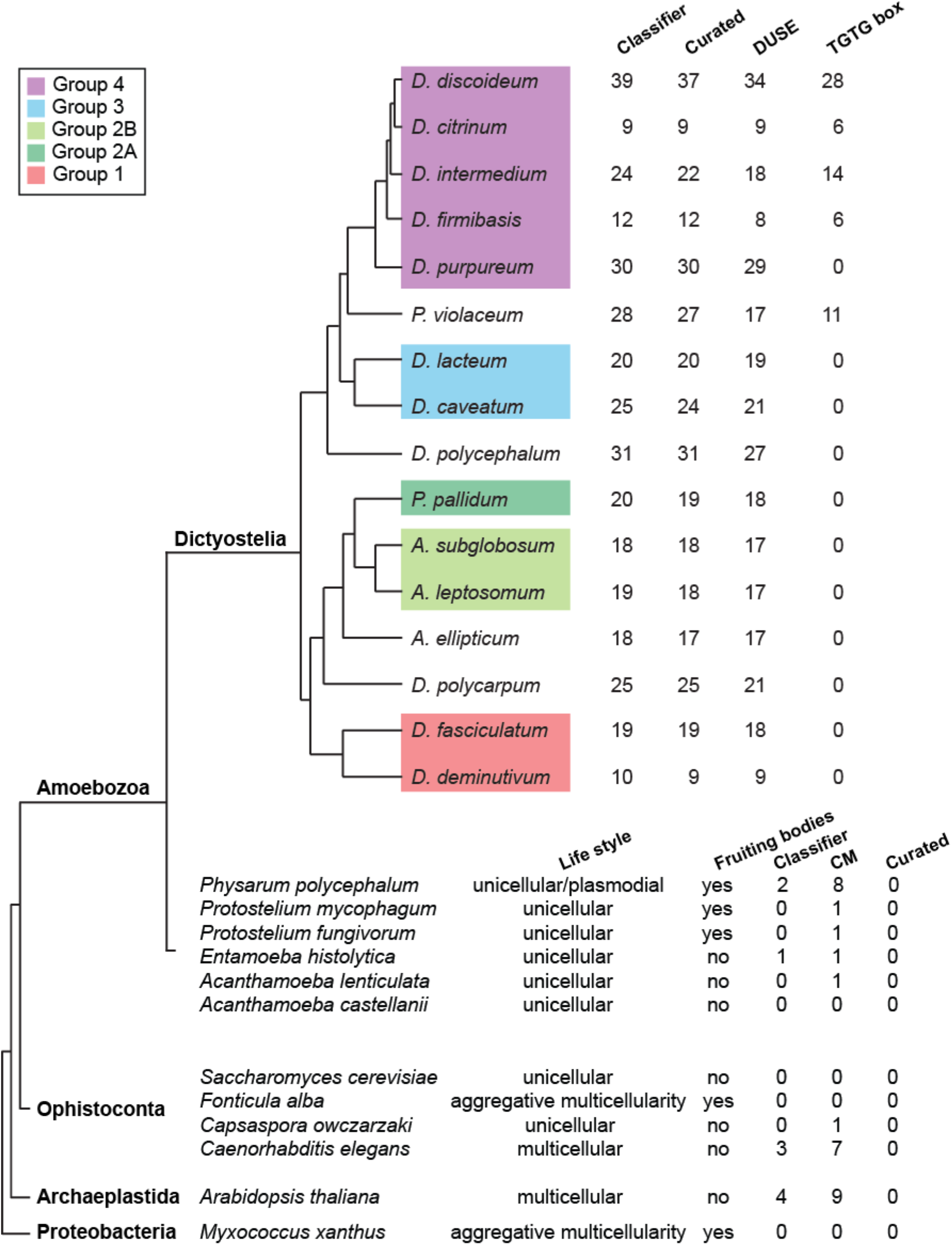
Class I RNAs are ubiquitous in and restricted to dictyostelid social amoebae. Upper part: Class I RNA loci were searched for in the genomes of 16 different Dictyostelia (Additional file 1). The number of hits identified by the classifier is indicated as well as the number of these loci that passed manual curation. The number of curated loci with DUSE and the TGTG-box at the correct distance are shown. Bottom part: Result from searches for Class I RNA loci outside Dictyostelia. Life style indicate unicellular or multicellular organisms. Fruiting bodies denotes if the organism life style involves formation of fruiting bodies. CM indicates number of candidates identified with a CM score ≥ 25. Headings Classifier and Curated as described above. Further information about outgroup Class I RNA candidates is available in Additional file 7.

### Genomic distribution of Class I RNA genes

In *D. discoideum*, all Class I RNA genes are located in intergenic regions and frequently found in clusters of two or more genes [33]. Clusters of at least two (different) Class I RNA genes are present in all analyzed Dictyostelia genomes, except for *D. citrinum* (Additional file 5). The absence of Class I RNA gene clusters in *D. citrinum* is likely a consequence of the quality of the genome assembly (Additional file 1), which is also reflected in the low number of identified Class I RNA genes. Clusters with a higher number of Class I RNA genes (three or more) are only found in five genomes, where the distribution in *P. pallidum* resembles that in *D. discoideum*, i.e. many of the genes are collected in two larger clusters on the same chromosome (Additional file 5). Even though many Class I RNA genes cluster together, they rarely have identical sequences. Only a few species-specific identical Class I RNAs were found in *D. polycephalum, P. violaceum, D. purpureum, P. pallidum* and *D. discoideum*, where in the latter two the identical genes are located in clusters (Additional file 5). Are there Class I RNAs that are identical between two different species? The only examples found were two Class I RNA loci shared between the group 4 species *D. discoideum* and *D. firmibasis* (Additional file 4).

To explore the origin of Class I RNAs further, we used the most well-annotated genomes to search for shared synteny by identifying orthologous genes in the 10 kb region flanking each Class I RNA locus (Materials and methods). Using this approach, we did not identify any strong evidence for shared synteny for Class I RNA genes between the different groups of Dictyostelia. Next we investigated if shared synteny could be detected within group 4 only by performing the same search using the genomes of *D. discoideum, D. purpureum* and *D. firmibasis*. Almost half of the Class I RNAs in *D. firmibasis* appear to share synteny with *D. discoideum* Class I RNAs, supported by several protein gene orthologues (Additional file 6) while no well-supported examples were found for *D. purpureum*. For the identical Class I RNA genes in *D. discoideum* and *D. firmibasis*, shared synteny was detected for DfiR-4 and DdR-47 (Additional file 6). Shared synteny between the other two identical Class I RNAs, DfiR-12 and DdR-50, could not be properly assessed due to the lack of available genome sequence surrounding DfiR-12.

### Class I RNAs are unique to dictyostelid social amoebae

The omnipresence of Class I RNAs within Dictyostelia, their developmental regulation as well as the aberrant development of *D. discoideum* cells lacking *DdR-21* [31] led us to hypothesize that this class of ncRNAs might be involved in the evolution of Dictyostelia aggregative multicellularity. In order to investigate this further, we searched for Class I RNA genes in genomes of unicellular amoebae and amoebae able to form unicellular fruiting bodies. Furthermore, we explored representative genomes of other major eukaryotic groups, i.e. archeaplastida and ophistokonta. We also chose to include the proteobacteria *Myxococcus xhantus* as these bacteria exhibit aggregative multicellularity [37], which in many aspects are analogous to Dictyostelia multicellularity (Fig. 4). We searched these genomes using the same successful approaches as for Dictyostelia, i.e. using the Class I classifier, based on promoter characteristics combined with RNA structure and sequence, as well as CM search alone. Only a few candidates were identified with the Class I classifier. This was anticipated since we did not expect the DUSE sequence or its distance to the TSS to be conserved outside Dictyostelia. The Infernal search resulted in a slightly higher number of Class I RNA gene candidates. However, manual inspection revealed that the candidates are unlikely to represent true Class I RNA genes as they were few in numbers and did not share characteristics, such as conserved 5’ and 3’ ends and presence of conserved sequence motif (Fig. 4, Additional file 7). Taken together, no Class I RNA genes were identified outside Dictyostelia, suggesting that this class of ncRNAs is unique to dictyostelid social amoebae and important for their aggregative multicellularity.

### Conserved Class I RNA interacting proteins

Class I RNAs are conserved throughout the evolution of Dictyostelia but does this also apply to proteins associated with this class of ncRNA? We previously identified four Class I RNA interacting proteins in *D. discoideum* by using one specific Class I RNA, DdR-21, as bait in pull-down experiments. Two of these proteins, GuaB and NdkC, are involved in nucleotide metabolism while the function of DDB_G0281243 is unknown. The fourth identified Class I RNA interacting protein, CIBP (also known as Rnp1A [38]), harbors two RNA binding motifs (RRMs) and was demonstrated to bind directly to the Class I RNA [31]. We searched for orthologues of the four proteins in the best annotated genomes of Dictyostelia (Fig. 1b, Additional file 1) and could identify orthologues to GuaB, DDB_G0281243 and CIBP in all Dictyostelia genomes investigated. Next, we used mRNA-seq data from *D. discoideum* (Group 4) [39] as well as *D. lacteum* (Group 3), *P. pallidum* (Group 2) and *D. fasciculatum* (Group 1) [20] to investigate expression and developmental regulation of the genes for each of the orthologues. According to the RNA-seq data, only *CIBP* were expressed in all four species. Interestingly, during early development the expression pattern of the gene is similar to that of Class I RNAs [31], i.e. *CIBP* appears to be down-regulated during early development in all species. However, as *D. discoideum* Class I RNAs continue to decrease in expression throughout development, *CIBP* expression seems to increase in the final stages of development (Additional file 2: Fig. S7a). Next, we performed gene predictions in the ten additional Dictyostelia genomes (Fig 4) and found *CIBP* orthologues in all species except for *D. citrinum* (Additional file 2: Fig. S7b). The absence of *CIBP* in *D. citrinum* is likely due to the quality of the genome assembly (Additional file 1). The majority of the orthologues are predicted to encode an approximately 300 amino acids (aa) long protein with two RRM’s. However, shorter orthologues were identified in *A. leptosomum* (114 aa), *A. ellipticum* (61 aa) and *D. purpureum* (203 aa). Also, only one RRM could be identified in the *CIBP* orthologues in *D. purpureum* and *A. ellipticum* (Additional file 2: Fig. S7b). Collectively, the majority of the disctyostelids are predicted to encode/express CIBP of similar length where the N- and C-terminal sequences contains RNA binding motifs, RRMs, while the central part of the protein is less conserved. Despite several efforts by us and others, all attempts to generate a CIBP knock out strain in *D. discoideum* have been unsuccessful [31, 38]. In addition, no CIBP mutants were generated in the recent Genome Wide *Dictyostelium* Insertion (GWDI) project (www.remi-seq.org). Taken together, this indicates that the CIBP protein is essential.

## Discussion

The development of high-throughput sequencing techniques has led to the discovery of numerous ncRNAs. In particular, it has facilitated the identification of small ncRNAs such as mi- and si- and piwi-interacting (pi)RNAs sized ~21-31 nt and long ncRNAs that can consist of up to several thousand nt. However, “mid-sized” ncRNA have largely been overlooked partly due to the size selection commonly carried out before sequencing to enrich for small RNAs and to avoid abundant RNAs such as rRNAs and tRNAs or fragment thereof. Here we used genome analyses in combination with expression validation to prove the existence of Class I RNAs in 16 dictyostelid social amoeba. This indicates that Class I RNAs were present in the last common ancestor of Dictyostelia, dating back at least 600 million years, and were involved in the transition from unicellular to multicellular life.

Class I RNAs play an important role in *D. discoideum*, as suggested by e.g. the large number of highly expressed genes and requirement for normal multicellular development [30, 31]. This class of ncRNAs was initially discovered by sequencing cDNA libraries of full-length RNA sized 50-150 nt [30]. In the same study, a putative promoter element, DUSE, was identified to be associated with Class I RNA genes. Later, two approaches to predict Class I RNAs were published. Fragrep, a tool which predict ncRNAs based on sequence motifs separated by a variable region, identified 45 Class I RNA candidate genes in *D. discoideum* of which 34 had been previously experimentally validated [40]. In *D. purpureum*, another group 4 dictyostelid, 26 Class I genes were predicted by searching for enriched 8-mers downstream of DUSE motifs in the genome sequence [18]. Hence, previous to the present study, Class I RNAs had only been identified in organisms belonging to one group of Dictyostelia. We now asked if Class I RNA genes also are present in other organisms, both within Dictyostelia characterized by aggregative multicellularity and outside this group of social amoebae.

In order to detect Class I RNAs, we first built a covariance model (CM) which predicts candidates based on both sequence and structure information of Class I RNAs. Based on the CM, we created a classifier which evaluates the identified candidates based on the presence of DUSE at the correct distance from TSS or from the TGTG-box (found upstream of many *D. discoideum* Class I RNA genes). Next, we confirmed expression of approximately 100 Class I RNAs, predicted by the classifier, by northern blot and/or RNA-seq. We were now confident that we could use the Class I classifier to accurately identify expressed Class I RNA genes also in other organisms. Based on this, we show that Class I RNAs are present in all dictyostelids with available genome sequences (total of 16). Having established the presence of Class I RNAs in all tested disctyostelids, we turned our focus on organisms outside of Dictyostelia. Outgroups were chosen to both represent organisms with a unicellular life style and those that go through different kinds of multicellular development. We used both the classifier and the CM search alone to search for candidates, the latter to avoid the constraint of having a DUSE element in front of the Class I RNA genes. Regardless of search approach, we did not identify Class I RNAs in any organism outside Dictyostelia, suggesting that these RNAs are restricted to dictyostelid social amoebae.

The ubiquitous presence of Class I RNA genes in Dictyostelia and absence in the closest related unicellular amoebae suggest that these RNAs were present in the last common ancestor of Dictyostelia and played a role in the transition from unicellular life to aggregative multicellularity. This is further supported by the developmental regulation of Class I RNAs and that at least one member, DdR-21, is required for normal development [30, 31]. In *D. discoideum*, transition from unicellular growth to multicellular development is associated with large transcriptional reprogramming of protein coding genes [23] and different cell types, i.e. prespore and prestalk cells, can be separated based on the transcriptional signatures of individual cells [41]. Interestingly, the majority of the protein coding genes that are essential for multicellular development in *D. discoideum* is also present in strictly unicellular amoebae [20]. Hence, it is likely that the ability to regulate how genes are expressed have been key in the evolution of multicellular development. Maybe Class I RNAs can rewire gene expression of genes present in unicellular organisms to create new networks adapted for development. We have no evidence that Class I RNA directly interacts with mRNA to regulate gene expression even though we are not excluding this mode of action. Another possibility is that Class I RNA regulate development by binding to proteins that directly or indirectly control development, maybe by acting as a molecular sponge, where specific proteins are sequestered by Class I RNAs. This could buffer the action of these proteins. Perhaps the observed down regulation of Class I RNAs during development lead to an increase in free active proteins important for multicellular development. CIBP would be a candidate for this kind of regulation since CIBP directly interacts with Class I RNAs, at least with the tested DdR21, in *D. discoideum* [31]. Support for CIBP as an important protein for development is its presence in all dictyostelids. We are currently attempting to decipher the function of Class I RNAs in *D. discoideum* by using a coupled RNA-seq and proteomics approach to investigate the involvement of Class I RNA in early multicellular development. The tendency to differentially regulate genes in order to create new functions is seen also in metazoan evolution, where an increase in regulatory miRNAs is correlated with increased organismal complexity [2]. Interestingly, *D. discoideum* is one of the few organisms outside animals and land plants were miRNAs have been identified [24–27]. If miRNAs, similar to Class I RNAs, are present in other dictyostelids is currently being investigated.

In Dictyostelia, the most complex multicellularity is found within group 4, exemplified by regulated proportions of specialized cell types and a migrating slug stage [3]. Interestingly, in analogy to miRNA expansion in complex animals, the number of expressed Class I RNA genes is correlated with Dictyostelia complexity where group four has the largest number of Class I RNA genes (Fig. 2b). In addition, RNA-seq data indicate that Class I RNAs are expressed at higher levels in group 4 (*D. discoideum*), further strengthening that Class I RNAs are involved in increased organismal complexity. This increased expression appears to be connected to the TGTG-box between DUSE and the TSS. So far, the TGTG-box have only been identified in Class I RNA loci in species belonging group 4 (except *D. purpureum*) and *P. violaceum* complex (Figs 3a and 4 and Additional file 3). Thus, this motif is likely a rather late addition in the evolution of Class I RNAs. It should be noted that it is currently not known if the second motif is actually a promoter element or the TSS of a longer precursor that is processed down to the mature RNA. In either case, the TGTG-box is associated with Class I RNA genes in social amoebae with higher levels of complexity as compared to other dictyostelids and appears to add another layer of regulation to Class I RNA expression. The emergence of the TGTG-box somewhere after the split of group 3 and 4 and its connection to increased phenotypic complexity seen in group 4 dictyostelids is somewhat analogous to changes in cis-regulatory elements, such as enhancers, and morphological evolution in animals [42].

The high number of novel Class I RNA loci identified in Dictyostelia enabled comparative studies which provides information on their key features. The short stem connecting the 5’ and 3’ ends of mature Class I RNA is conserved. Furthermore, the sequence variability of the stem between organisms but also the high number of compensatory mutations within each species strongly suggest that it is the structure rather than sequence that is important for function. Flanking the stem structure at both ends are highly conserved nucleotides present in almost all Class I RNAs: AAG and CTGT at the 5’ and 3’ends respectively, where the 5’G and 3’C forms the first base-pair of the stem. Curiously, in spite of the high conservation of the three 5’ nucleotides, the mature transcript of Class I RNAs in *D. discoideum* almost always starts with the G residue, while it is more common in the group 1-3 representatives that also the two A’s are included. In either case, the start of each mature Class I RNA is well defined with the great majority of all RNA-seq reads sharing the same 5’ end. The 3’end is also remarkably conserved. Whether these nucleotides are part of the mature transcript is hard to assess due to the sequencing approach used, which only allowed us to capture a small fraction of the 3’ ends. However, we do find reads covering the 3’ conserved CTG in the majority of the different Class I RNAs, suggesting that these constitute the 3’ end of the mature Class I RNAs. This is also supported by the full-length cDNA libraries of small RNAs in *D. discoideum* where this class of RNA was first discovered [30].

The comparison of Class I RNA loci also confirms the importance of the ~11 nt motif present immediately after the 5’ part of the stem structure. Although some of the nucleotides vary within and in between organisms, several residues are nearly perfectly conserved. Yet another conserved feature is the putative promoter motif, DUSE. Based on studies of spliceosomal RNAs in *D. discoideum*, we previously demonstrated that DUSE is associated with genes transcribed by both RNA Pol II and RNA Pol III [43]. However, we believe that Class I RNAs are transcribed by RNA Pol III since the canonical RNA Pol III termination signal is present downstream of most Class I RNA loci and Class I RNA reads are lacking in poly(A) enriched RNA-seq libraries.

Based on the findings in this study, we conclude that Class I RNAs were present in the last common ancestor of Dictyostelia. In addition, our data also strongly suggest that the putative promoter element DUSE was present 60 nt upstream of the ancestral Class I RNA gene and that the element was required for expression of the gene. We also conclude that the ancient Class I RNA was characterized by a short stem structure and a 11 nt sequence motif where at least six of the positions were identical to the corresponding nucleotides in extant Class I RNAs. Due to the high sequence and structure variability of the region between the 11 nt motif and the start of the 3’ stem in identified Class I RNAs, we cannot resolve this part of the ancestral sequence. However, the total length of the mature RNA was probably approximately 60 nt long (Fig. 5).

**Figure 5.**
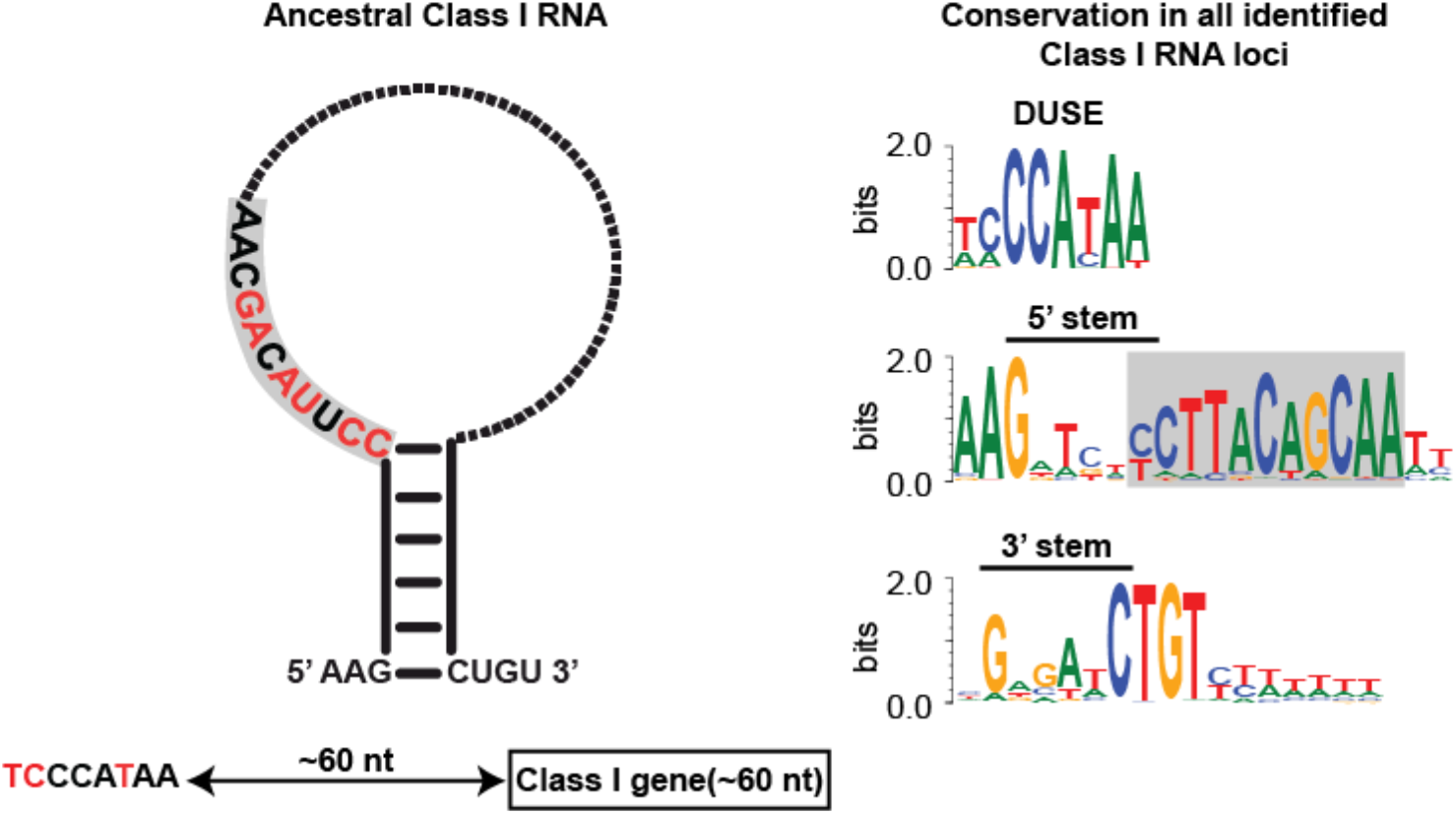
Ancestral Class I RNA and conserved key features. Left: Schematic representation of the ancestral Class I RNA transcript. The putative promoter element DUSE and its distance to the Class I RNA gene is indicated below the Class I RNA structure. Strongly conserved features, nucleotide and base paired stem, are colored black while red denotes more variable positions (based on data presented in figure 3). The dotted part of the loop indicates highly variable sequence. Right: Sequence logos of alignments of conserved sequence motifs from all identified and curated Class I RNAs. Only DUSE identified at the correct distance were included in the alignment. The 11 nt motif is boxed in gray in both the ancestral Class I RNA (left) and sequence logo (right).

Even though some motifs are highly conserved within all Class I RNAs, conservation of complete Class I RNA genes are rare. Only a few examples of identical loci within the same genome are found in a handful of dictyostelids and only two identical Class I RNAs shared by two different species were identified, i.e. the group 4 dictyostelids *D. discoideum* and *D. firmibasis* (Additional file 4). In *D. discoideum* and *P. pallidum* many Class I RNA genes are situated in larger clusters and it is within these clusters where the species-specific identical Class I RNA genes are found, perhaps indicating expansion of Class I RNA genes by duplication. Interestingly, the snRNA genes in *D. discoideum* are organized in a similar way where closely related genes often are found in pairs situated very close together [33]. However, in general the Class I RNA genes are spread out in the genomes and this is also true for the majority of the identical Class I RNA genes in the different genomes. Hence, the low occurrence of shared synteny, low overall sequence conservation and different number of loci in different organisms suggest that the expansion of Class I RNAs mainly occurred after speciation.

## Conclusions

We have identified Class I RNAs in 16 different dictyostelids and validated their expression in representatives of each major group of Dictyostelia dating back at least ~600 million years. Despite the large evolutionary distances, Class I RNA genes share promotor motifs and the mature RNAs have several characteristics in common, i.e. short stem, conserved sequence motif and highly conserved 5’ and 3’ ends. In addition, the *D. discoideum* Class I RNA interacting protein CIBP is conserved throughout Dictyostelia. Furthermore, the gene shares its expression profile with Class I RNAs during early development, indicating that function and mode of action is also conserved. Although Class I RNAs are present in all dictyostelids investigated, no evidence was found for this class of RNAs in any other organism. Taken together, our results suggest that Class I RNAs are important for the evolution of multicellularity in Dictyostelia.

## Materials and methods

### RNA isolation and northern blot

The following strains were used for northern blot validation of Class I expression: *D. discoideum* AX2 (DBS0235521, www.dictybase.org), *D. purpureum* WS321, *P. pallidum* PN500, *D. lacteum* Konijn, *D. fasciculatum* SH3, *A. subglobosum* LB1. All strains, except *D. discoideum*, were kindly provided by Dr Maria Romeralo and Professor Sandra Baldauf. Total RNA was extracted with TRIzol (ThermoFisher Scientific) from cells grown in association with *Klebsiella aerogenes* on non-nutrient agar. Northern blots were performed as described in [30, 31]. Briefly, 10 μg total RNA was separated on 8 % PAGE/7 M Urea and electroblotted to Hybond N+ nylon membranes (GE Healthcare). After UV crosslinking, immobilized RNA was hybridized with ^32^P-labeled oligonucleotides in Church buffer at 42 °C overnight. Signals were analyzed with a Personal Molecular Imager (BIO-RAD) normally after a few hours’ exposure. Membranes analyzed more than once were stripped with 0.1×SSC/1 % SDS buffer at 95 °C for 1 h and controlled for residual signal before reprobing. Oligonucleotide sequences are provided in Additional file 2: Table S1.

### RNA-seq validation

Strain growth and RNA extraction for RNA-seq have been described previously [20]. For each strain, RNA was prepared from growing cells and two multicellular developmental stages: aggregates and tipped aggregates/fingers (biological duplicates except for *D. lacteum*). Truseq small RNA Sample Preparation kit (product # RS-200-0012, Illumina) was used to prepare sequencing libraries from 1 μg total RNA. The library preparation was performed according to the manufacturer’s protocol (#15004197 rev G) where cDNA representing 18 nt – 70 nt RNA were isolated. Single read 50 bp sequencing was performed using v4 sequencing chemistry on an Illumina HiSeq2500. To reduce influence of degradation products, only full length 50 bp reads were mapped with bowtie allowing for 1 mismatch [44] and counted with feature counts [45]. Read coverage over Class I candidate loci were calculated with BEDTools genomecov v. 2.26.0 [46]. Class I RNAs were considered to be validated by RNA-seq if the read coverage indicated a distinct 5’ end and reads specifically matched the predicted loci, i.e. did not appear to be part of a considerably longer transcript. Principal component analyses were performed using DESeq2 [47]. ROC plot evaluation of Class I prediction was performed using pROC package[48] in R.

### Strains and genomic resources

Strain names and accession numbers for genomic sequences are listed in Additional file 1.

### Identification of Class I RNAs

Of the 40 *D. discoideum* Class I RNAs annotated previous to this study, six were excluded from the model build (r48, r53, r54, r55, r58 and r61) as they represented truncated fragments or lacked the canonical features such as the stem or conserved sequence motif. The remaining 34 were aligned with MAFFT v7.407 [49] using the ginsi setting. Consensus structure for the alignment was predicted with RNAalifold 2.3.3 [50] using the -T 22 option to account for the optimal growing temperature of the amoebae. Alignment and consensus structure was combined to a Stockholm alignment file and a co-variance model was created with Infernal 1.1.2 [32]. Infernal was then used to search the genomes of *D. discoideum, D. purpureum, D. lacteum, P. pallidum, A. subglobosum* and *D. fasciculatum* using default settings and candidates with a score ≥ 25 were added to the alignment using MAFFT (ginsi -add). The new alignment was manually curated and used to predict consensus structure, create a new co-variance model, and perform a new search in the same genomes. This procedure was iterated six time, i.e. until no new candidates with an Infernal score ≥ 25 were identified. Enriched sequence motifs were identified up to 150 nt upstream of identified candidates with MEME v. 5.0.3 [51]. For final Class I identification, Infernal searches were performed with increased sensitivity and all candidates scoring 15 or higher were kept (cmsearch –nohmm –notrunc -T 15). The candidates were then evaluated based on the presence of DUSE and TGTG-box in the 150 nt preceding the predicted start of transcription (see above) using FIMO v. 5.0.3 [34]. Infernal score, FIMO motif score, and a motif distance score (+5) were then added to a total score. Missing DUSE or incorrect distance was penalized with −10 or −5 respectively. If a total score of 32 was achieved, the candidate was considered likely to be an expressed true Class I RNA and kept for further analyses. Representative sequence logos of manually curated sequence alignments (mafft --maxiterate 1000 –localpair) were created with WebLogo 3 [52].

### Orthologue identification and shared synteny search

For Dictyostelia species lacking genome annotations, gene prediction was performed with Gene id v. 1.4 [53] using Dictyostelium parameter file. Orthologue identification was performed using OrthoFinder v. 2.3.3 [54]. Protein domain architectures for CIBP/Rnp1A orthologues were analyzed with hmmscan using the HMMER web interface [55]. Orthologue information for *D. discoideum, D. firmibasis, D. lacteum, P. pallidum, A. subglobosum* and *D. fasciculatum* was used to investigate shared synteny for Class I RNA loci. For each Class I RNA locus, gene information within a 10 kb flanking region was retrieved. Next, we searched for orthologues of these genes in the other organisms included in the search. If a Class I RNA was found within 10 kb of an orthologous gene in another organism, it was manually inspected to determine the level of shared synteny.

## Declarations

### Ethics approval and consent to participate

Not applicable

### Consent for publication

Not applicable

### Availability of data and material

#### Competing interest

The authors declare that they have no competing interests

#### Authors’ contribution

JK, LA, JR, and FS participated in the design of the project; LA and ZL generated experimental data; JK performed most of the data analysis and prepared figures; JR participated in the bioinformatic analysis; L.E, A.N, G.G, and P.S supplied RNA for sequencing. JK and FS drafted the manuscript.

#### Funding

This work was supported by the Linneus Support from the Swedish Research Council to the Uppsala RNA Research Centre; Carl Tryggers Foundation under Grant number CST 18:381.

## Acknowledgements

Sequencing was performed by the SNP&SEQ Technology Platform in Uppsala. The facility is part of the National Genomics Infrastructure (NGI) Sweden and Science for Life Laboratory. The SNP&SEQ Platform is also supported by the Swedish Research Council and the Knut and Alice Wallenberg Foundation.

## Description of additional files

**File name:** Additional file 1

**File format:** excel (.xlsx)

**Title of data:** Genomic resources used in this study.

**Description:** References for dictyostelid genomic sequences used in the study together with alternative names (Synonyms) for organisms (where applicable) and basic genome sequence statistics. Genome sequences used in the construction of the co-variance model (CM) are indicated in bold. For outgroup genome sequences used in the study, only the GenBank or RefSeq reference are given.

**File name:** Additional file 2

**File format:** PDF (.pdf)

**Title of data:** Supplementary figures and tables

**Description:** Supplementary figures 1-7 and table 1

**File name:** Additional file 3

**File format:** excel (.xlsx)

**Title of data:** Spreadsheet with search summary and expression validation for Class I RNAs from selected dictyostelids.

**Description:** The spreadsheet show data for all predicted Class I RNAs, from respective organism, passing the CM score ≥ 15. Candidates considered to be true Class I RNAs have been named according to previous naming rules. Classifier summary includes classifier score and infernal (CM) score as well as identified DUSE sequence (DUSE_seq), the duse score obtained from FIMO (duse_score), the duse distance from the predicted Class I RNA start (duse_dist), the TGTG-box sequence (TGTG-box_seq) and score obtained from FIMO (TGT-box_score), TGTG distance from predicted Class I start (TGT-box_dist) and the distance between DUSE and TGTG-box (inter_dist). Read counts denotes raw RNA-seq read counts for each Class I RNA candidate in each sequencing library for the organisms where we performed RNA-seq. Read counts include time points, for growning cells (0h) and time after starvation, when RNA was collected and A and B denotes biological replicates. Expression validation is based on read counts and coverage over each respective locus and northern blot. For Read coverage, the values 1 and 0 denotes validated and not validated, respectively. The read coverage validation in combination with either CM score or classifier score was used to generate ROC curves in figure 2 and Additional file 2: Fig. S3. Northern blot indicates Class I RNAs analyzed by northern blot in this study.

**File name:** Additional file 4

**File format:** excel (.xlsx)

**Title of data:** Spreadsheet with name, genome location, validation and score of all curated Class I genes for all investigated dictyostelids.

**Description:** For each Class I RNA gene Genomic location includes chromosome/contig/scaffold (Chromosome), start and end of predicted RNA (Start and Stop, respectively), on which DNA strand the RNA is located (Strand), and the predicted length (Class I RNA length). The Classifier summary includes the classifier score, the CM score the DUSE sequence and score (DUSE_seq and duse_score, respectively)) and TGTG-box sequence and score (TGTG-box_seq and TGTG-box_score, respecively). Columns duse_dist and TGTG-box_dist denotes the distance between the motif and predicted TSS and inter_dist denotes the distance between the two motifs. Under Upstream motifs, columns DUSE and TGTG-box summarize the classifier output and indicate if respective motif is found at the correct distance (1) or not (0). Where applicable, information about RNA-seq (1 denotes expressed Class I based on read coverage and 0 if expression could not be validated) and/or northern blot validation is given.

**File name:** Additional file 5

**File format:** PDF (.pdf)

**Title of data:** Genomic distribution of Class I RNA loci.

**Description:** Chromosome/contig/scaffold names are indicated to the left and respective lengths are normalized to the longest one (length in bp are presented to the very right of each schematic stretch of DNA). Total number of Class I RNA genes for each organism is given at the top and for each chromosome/contig/scaffold to the very right. Class I RNA genes are indicated by black arrows except for genes with identical sequences, which are indicated with colored arrows. Short vertical lines specify every 0.5 mbp. In *D. discoideum*, the duplication on chromosome 2 (DDB0232429) is indicated by grey boxes with vertical lines.

**File name:** Additional file 6

**File format:** PDF (.pdf)

**Title of data:** Shared synteny between *D. discoideum* and *D. firmibasis* Class I RNA loci.

**Description:** Representation of *D. firmibasis* (dfi) Class I RNA loci +/- 10 kb that share synteny with *D. discoideum* (ddi) Class I RNAs supported by at least two orthologous genes (connected with dashed lines). Chromosome/contig names are given to the left of the genes. Genes situated on the forward and the reverse strand are indicated above and below the chromosome/contig, respectively. Vertical grey lines are given at every 1000 nt.

**File name:** Additional file 7

**File format:** excel (.xlsx)

**Title of data:** Identified Class I RNA candidates in outgroups.

**Description:** For each candidate (classifier score ≥ 32 and/or CM score ≥ 25) the following information is given: classifier score, CM score as well as identified DUSE sequence (DUSE_seq), the duse_score obtained from FIMO, the distance of DUSE (duse_dist) from the predicted Class I start, the TGTG-box sequence (TGTG-box_seq) and score (TGTG-box_score) obtained from FIMO, TGTG_distance from predicted Class I start and the distance between DUSE and TGTG-box (inter_dist). The last column (Seq), contains the full genomic sequence of each candidate.

## Notes

### Competing Interest Statement

The authors have declared no competing interest.

