## Additional file 2 for "Abundantly expressed class of non-coding RNAs conserved through the multicellular evolution of dictyostelid social amoebae"

### Additional file 2 – Supplementary figures and tables

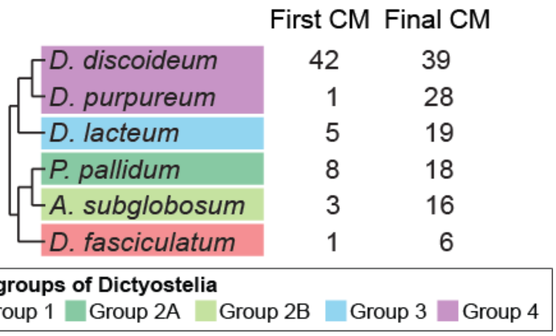


**Figure S1. Summary of the initial search (CM) for Class I RNA candidates.** Distribution of Class I RNA gene candidates (CM score ≥ 25) identified with the first version of the co-variance model (First CM) and the last version of the CM (Final CM). The first version of the CM was based solely on previously published *D. discoideum* Class I RNAs, while the final version of the CM was based on Class I RNA gene candidates from all six dictyostelids included in the CM build. The final CM search was performed with increased sensitivity (see Material and methods for details).

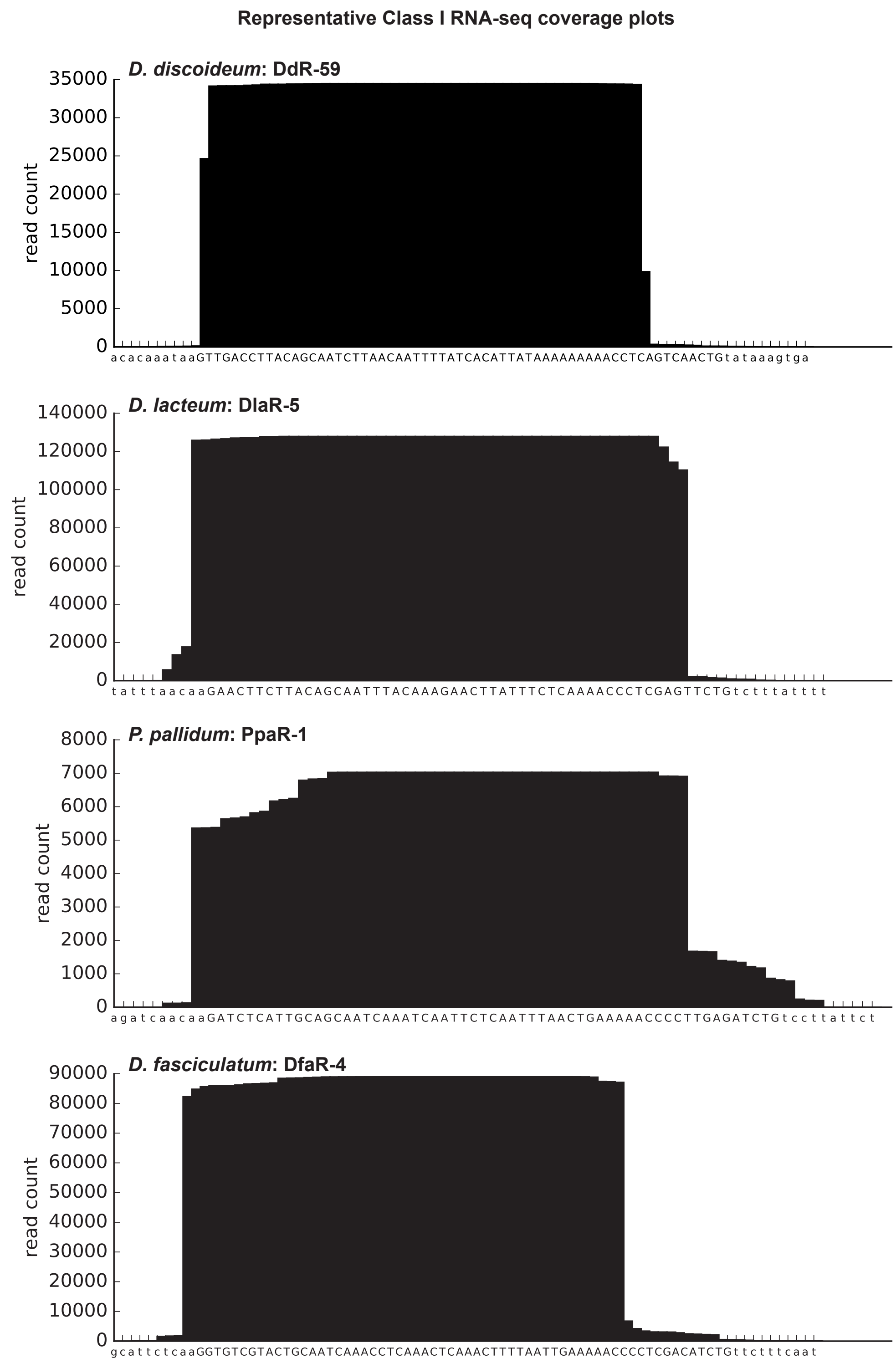


**Figure S2. Representative RNA-seq coverage plots of Class I RNAs.** Each Class I RNA plot represent the total raw read coverage for all replicates combined. Capital letters (x-axis) denote the nucleotide sequence of the predicted Class I RNA sequence and letters in lower case represent 10 nucleotides upstream and downstream of the 5’ and 3’ ends. DlaR-5, PpaR-1 and DfaR-4 was also validated by northern blot (see Fig. 2).

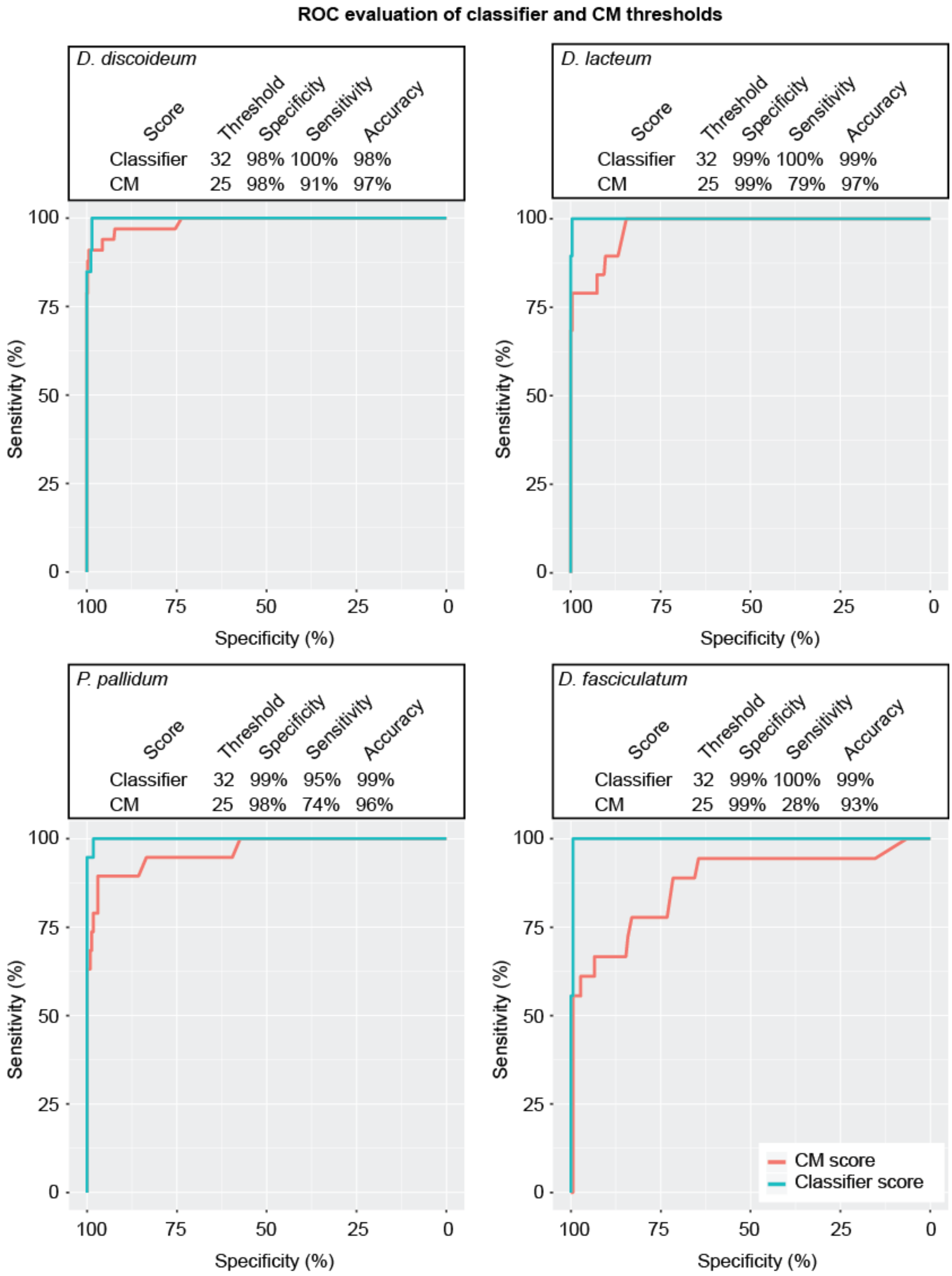


**Figure S3. Per species ROC evaluation of CM and Classifier thresholds.** Curves are based on the RNA-seq validation of all Class I RNA candidates (CM score ≥ 15) and either classifier score or CM score. The specificity, sensitivity and accuracy at the thresholds used for the classifier (32) and CM (25) is indicated above each plot. The input data used to generate the plots can be found in Additional file 3.

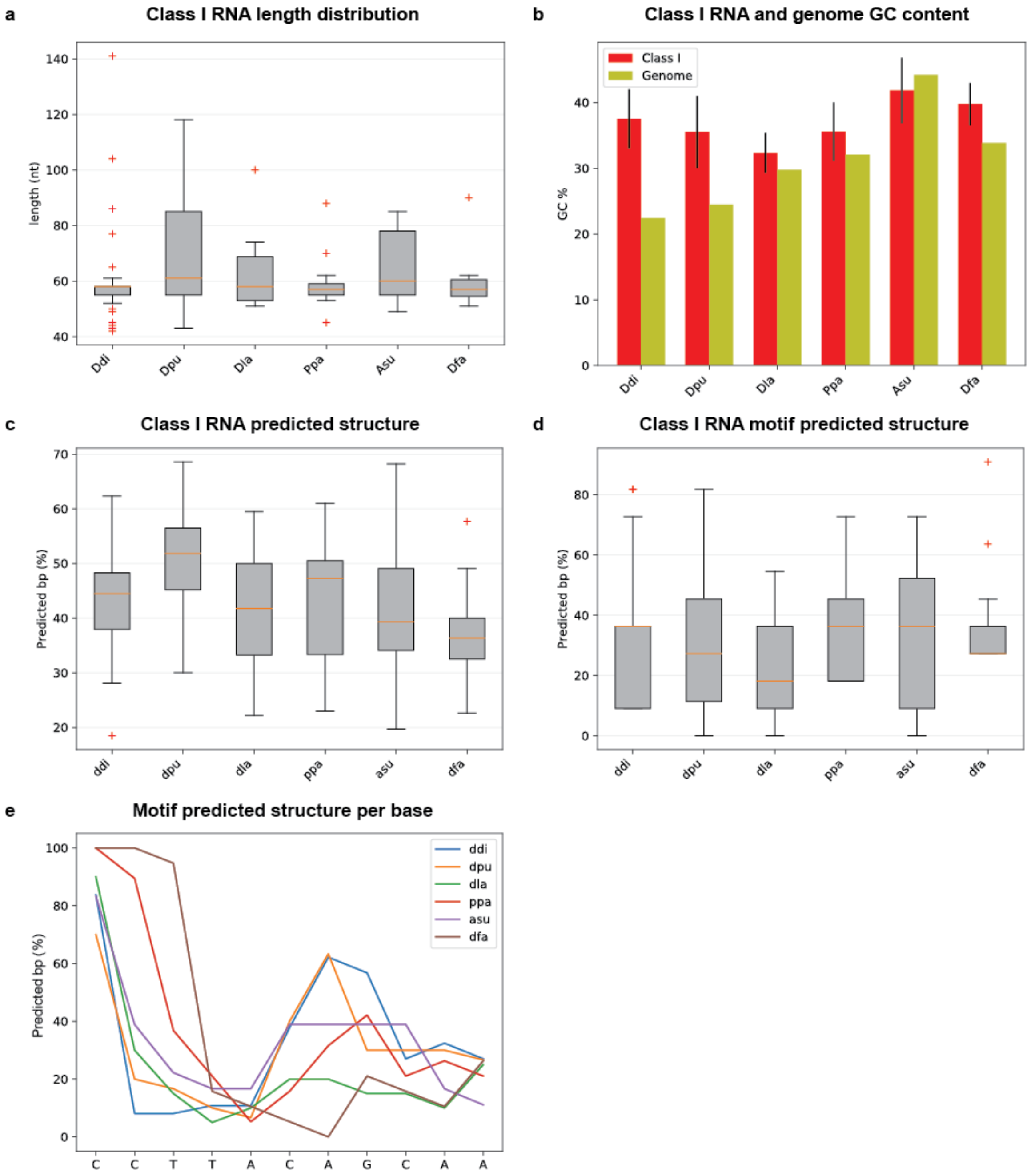


**Figure S4. Conserved features of Dictyostelia Class I RNAs in representative species.** a,c,d: boxes represent the quartiles of the lengths, whiskers represent the rest of the distribution except for outliers indicated with +. Orange solid line in boxes indicate the median. a) Length distribution of all identified Class I RNAs per species. b) Mean GC content (%) of Class I RNA and respective genome sequence. Error bars indicate standard deviation. c) Percent predicted base pairs of mature Class I RNAs. d) Percent predicted base pairing in the conserved 11 nt motif region. e) Predicted base pairing frequency per nucleotide for the conserved 11 nt motif. *D. discoideum* motif sequence is given on the x-axis and y-axis denotes percent of Class I RNAs that are predicted to be structured at the respective position of the motif. NB that the nucleotides are from the *D. discoideum* motif.

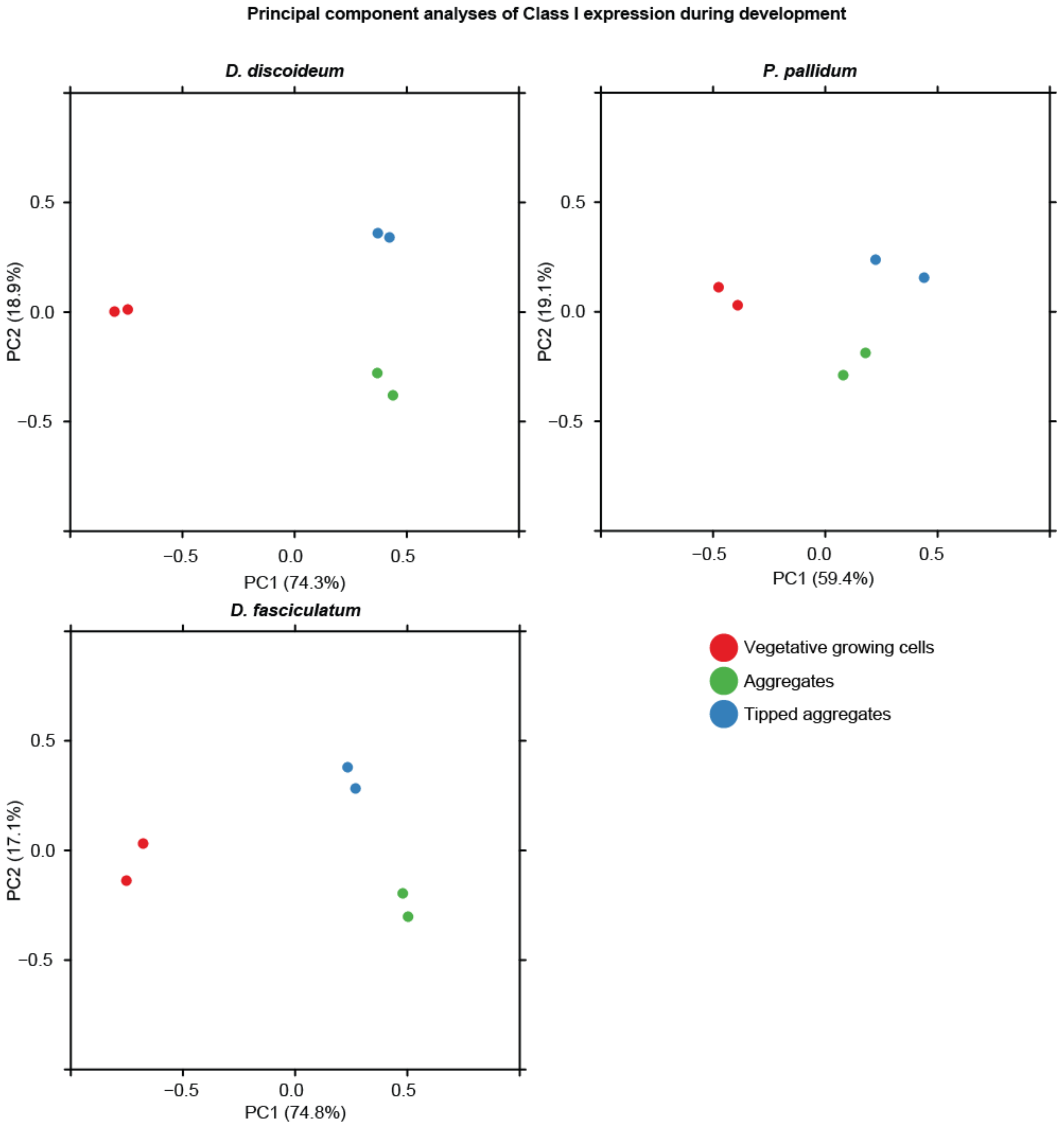


**Figure S5. Class I RNAs are developmentally regulated.** Principal component analysis based on RNA-seq read counts for Class I RNAs in *D. discoideum, P. pallidum* and *D. fasciculatum.* Developmental time points/structures (biological duplicates) are indicated by different colors.

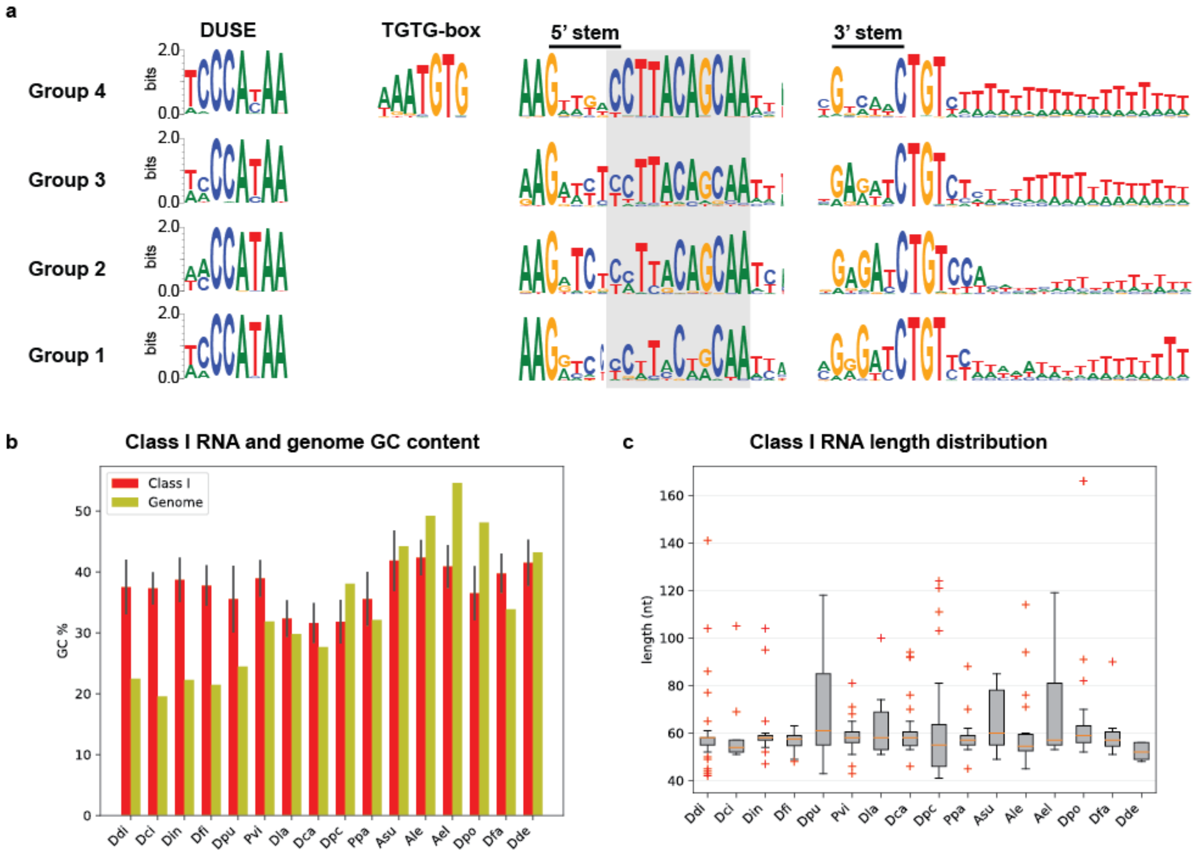


**Figure S6. Conserved features identified in representative species are conserved throughout Dictyostelia.** a) Conservation of DUSE, TGTG-box, 5' and 3' stem, motif sequence (boxed in gray) and downstream region for all Class I RNAs in each major subgroup of Dictyostelia. Two *D. citrinum* Class I RNAs were excluded from the alignments due to the lack of flanking genome sequences. b) Mean GC content (%) of Class I RNAs and respective genome sequence. Error bars indicate standard deviation. c) Length distribution of all identified Class I RNAs per species. Boxes represent the quartiles of the lengths, whiskers represent the rest of the distribution except for outliers indicated with +. Orange solid line in boxes indicate the median.

a

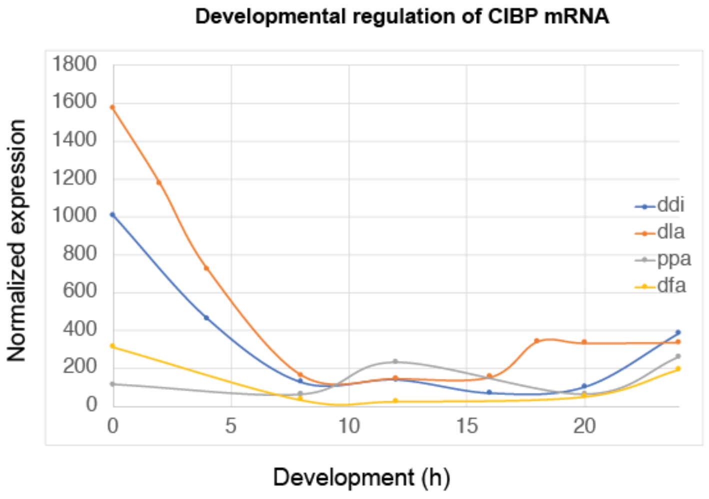


b

| | CIBP orthologues | RRM's per orthologue | Length |
|---|---|---|---|
| *D. citrinum* | 0 | n/a | n/a |
| *D. intermedium* | 1 | 2 | 297 aa |
| *D. firmibasis* | 1 | 2 | 295 aa |
| *D. purpureum* | 1 | 1 | 203 aa |
| *P. violaceum* | 1 | 2 | 273 aa |
| *D. lacteum* | 1 | 2 | 302 aa |
| *D. caveatum* | 1 | 2 | 288 aa |
| *D. polycephalum* | 1 | 2 | 289 aa |
| *P. pallidum* | 1 | 2 | 261 aa |
| *A. subglobosum* | 2 | 2 | 271/275 aa |
| *A. leptosomum* | 1 | 2 | 114 aa |
| *A. elipticum* | 1 | 1 | 61 aa |
| *D. polycarpum* | 1 | 2 | 274 aa |
| *D. fasciculatum* | 1 | 2 | 323 aa |
| *D. deminutivum* | 2 | 2 | 278/282 aa |

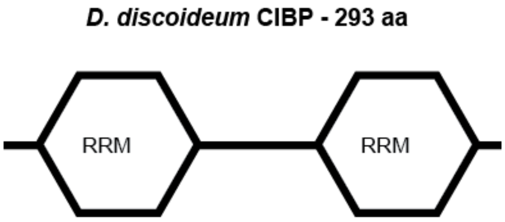


**Figure S7. Conservation of the Class I RNA interacting protein CIBP in Dictyostelia. a)** Developmental regulation of CIBP mRNA in *D. discoideum* (ddi), *D. lacteum* (dla), *P. pallidum* (ppa), and *D. fasciculatum* (dfa). Developmental time points are normalized to the development of *D. discoideum* (Expression data from [1, 2]. b) Number of CIBP orthologues identified in each species, number of RNA recognition motif's (RRM's) per orthologue, and the predicted length of the protein. The domain composition of *D. discoideum* CIBP, which is representative for all CIBP with two RRM's, is schematically depicted to the right.

**Table S1.** Oligonucleotides used for northern blot analysis. Name, sequence and matching Class I RNA(s) are denoted for each oligonucleotide/probe.

| Oligo name | Oligo sequence | Class I RNA |
|---|---|---|
| DfaR-4 | GTTTGATTGCAGTACGACACC | DfaR-4 |
| AsuR-10 | GTAGGATACGTTTGTTTGGTTG | AsuR-10 |
| AsuR-13/14 | GTTTGAAGTTTGGTAGTGATT | AsuR-13, AsuR-14 |
| PpaR-9 | GTTGATTGCTGTAAGGCCACC | PpaR-9 |
| PpaR-1/2 | GGGTTTTTCAGTTAAATTGAG | PpaR-1, PpaR-2 |
| DlaR-5 | GAGAAATAAGTTCTTTGTAAA | DlaR-5 |
| DlaR-1 | GGAGAAAAAAGATCATTTAG | DlaR-1 |
| DpuR-X | CCAATTTTTCCCGGAAAGAC | DpuR-5, DpuR-6, DpuR-13, DpuR-16, DpuR-21, DpuR-26 |
| DpuR-7 | CTTTGGGAGGGAAAATGTCAA | DpuR-7 |
| DpuR-Y | TTGCTGTAAGGAATTCTTGA | DpuR-2, DpuR-10, DpuR-12, DpuR-6, DpuR-24, DpuR-1, DpuR-26, DpuR-21, DpuR-7, DpuR-8, DpuR-4, DpuR-3, DpuR-17, DpuR-13, DpuR-23, DpuR-18, DpuR-20, DpuR-9, DpuR-16, DpuR-5, DpuR-19, DpuR-14, DpuR-11, DpuR-25 |
