## Additional file 5 for "Abundantly expressed class of non-coding RNAs conserved through the multicellular evolution of dictyostelid social amoebae"

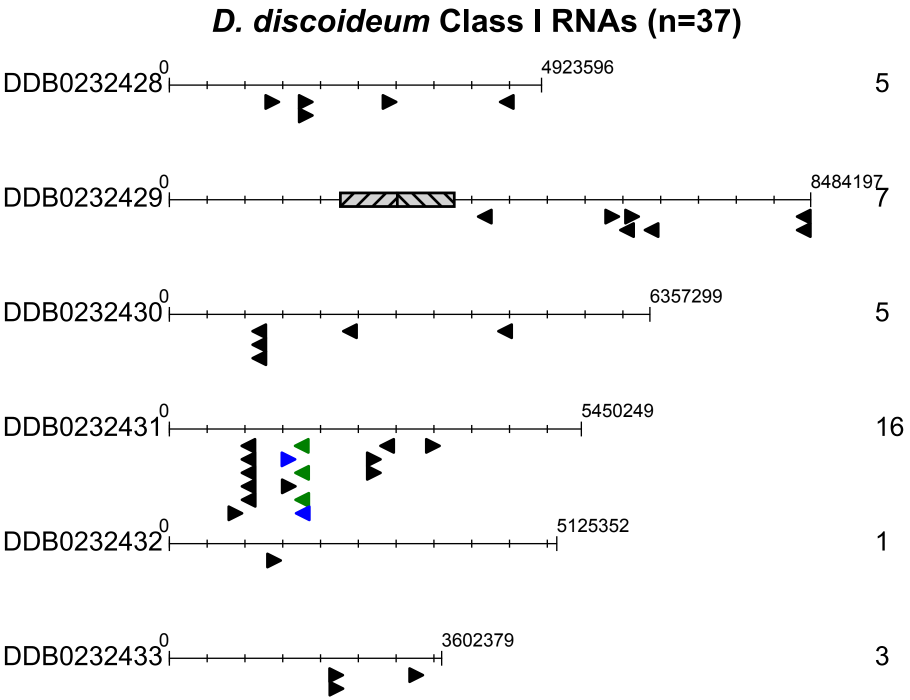
D. discoideum Class I RNAs (n=37)
DDB0232428
0
4923596
5
DDB0232429
0
8484197
7
DDB0232430
0
6357299
5
DDB0232431
0
5450249
16
DDB0232432
0
5125352
1
DDB0232433
0
3602379
3

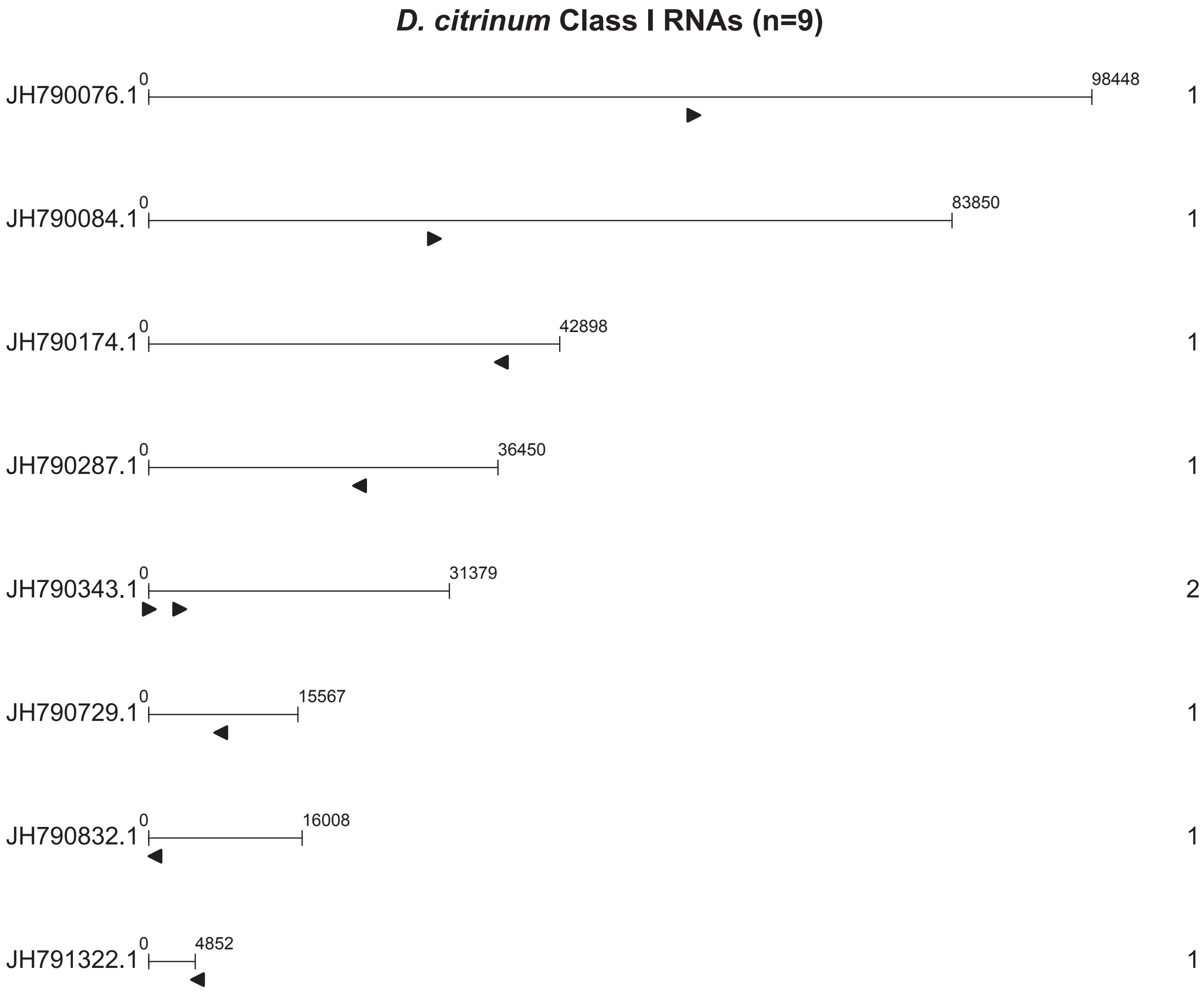
D. citrinum Class I RNAs (n=9)
JH790076.1
0
98448
1
JH790084.1
0
83850
1
JH790174.1
0
42898
1
JH790287.1
0
36450
1
JH790343.1
0
31379
2
JH790729.1
0
15567
1
JH790832.1
0
16008
1
JH791322.1
0
4852
1

***D. intermedium* Class I RNAs (n=22)**

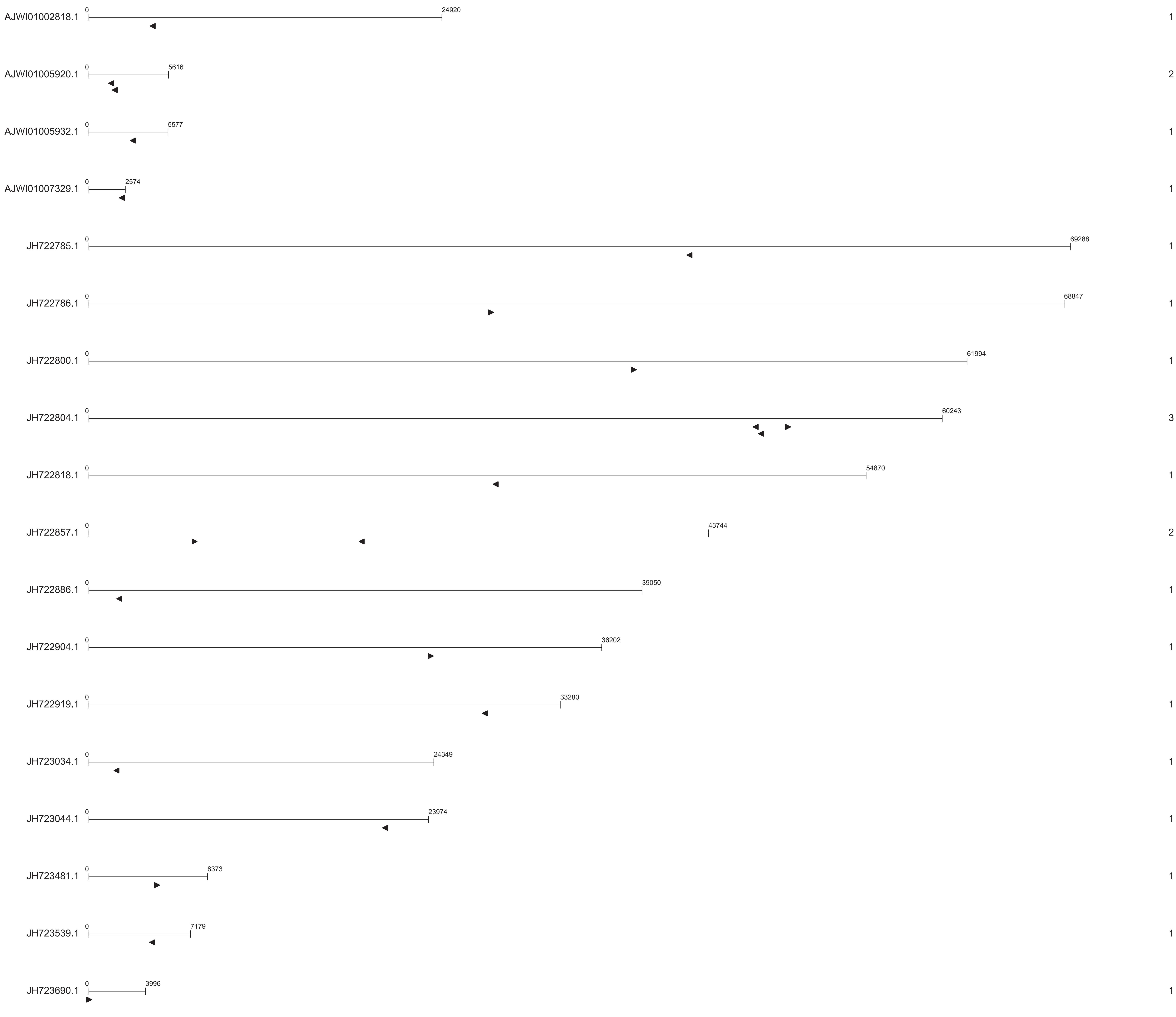

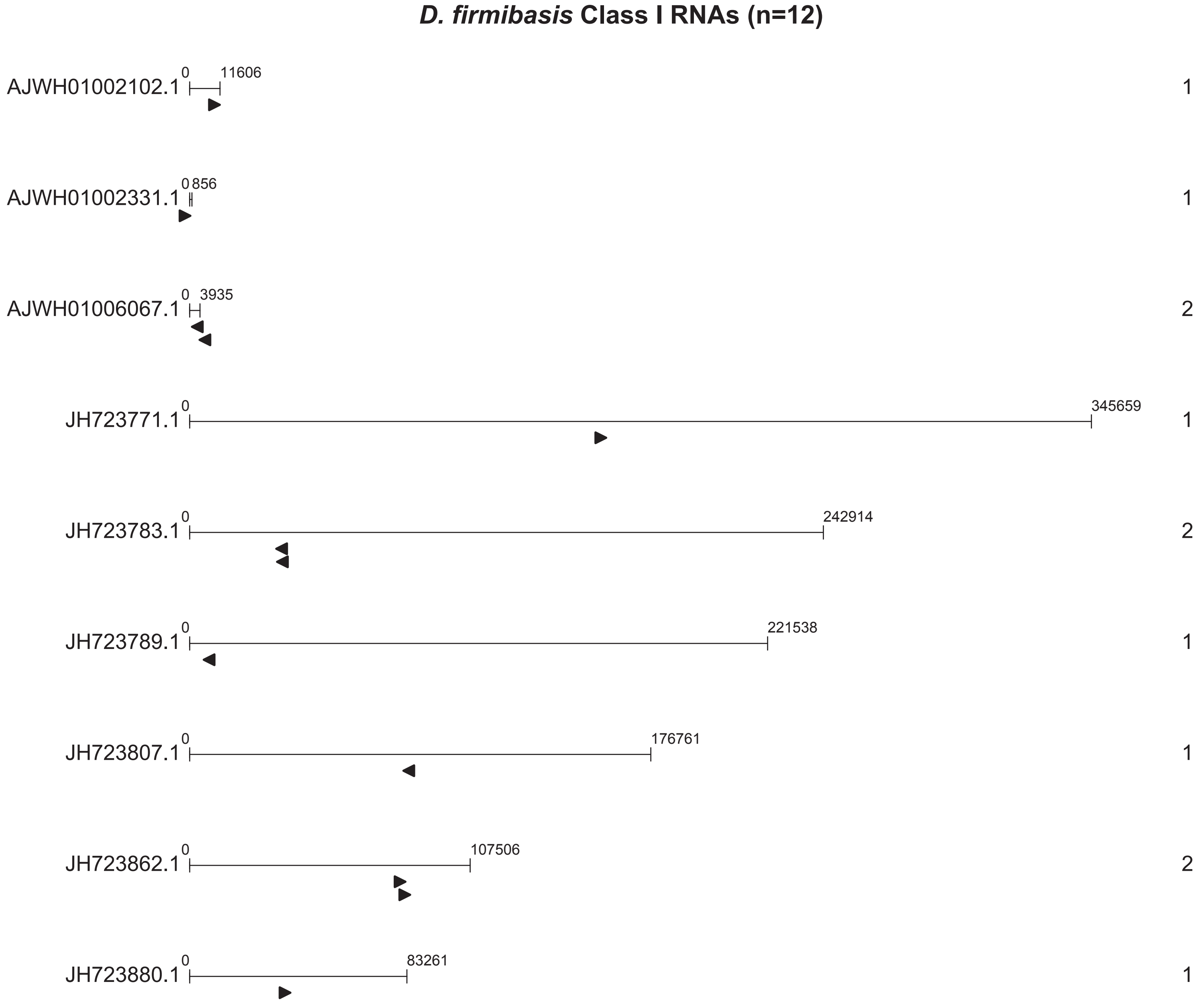
D. firmibasis Class I RNAs (n=12)
AJWH01002102.1
0
11606
1
AJWH01002331.1
0
856
1
AJWH01006067.1
0
3935
2
JH723771.1
0
345659
1
JH723783.1
0
242914
2
JH723789.1
0
221538
1
JH723807.1
0
176761
1
JH723862.1
0
107506
2
JH723880.1
0
83261
1

***D. purpureum*** **Class I RNAs (n=30)**

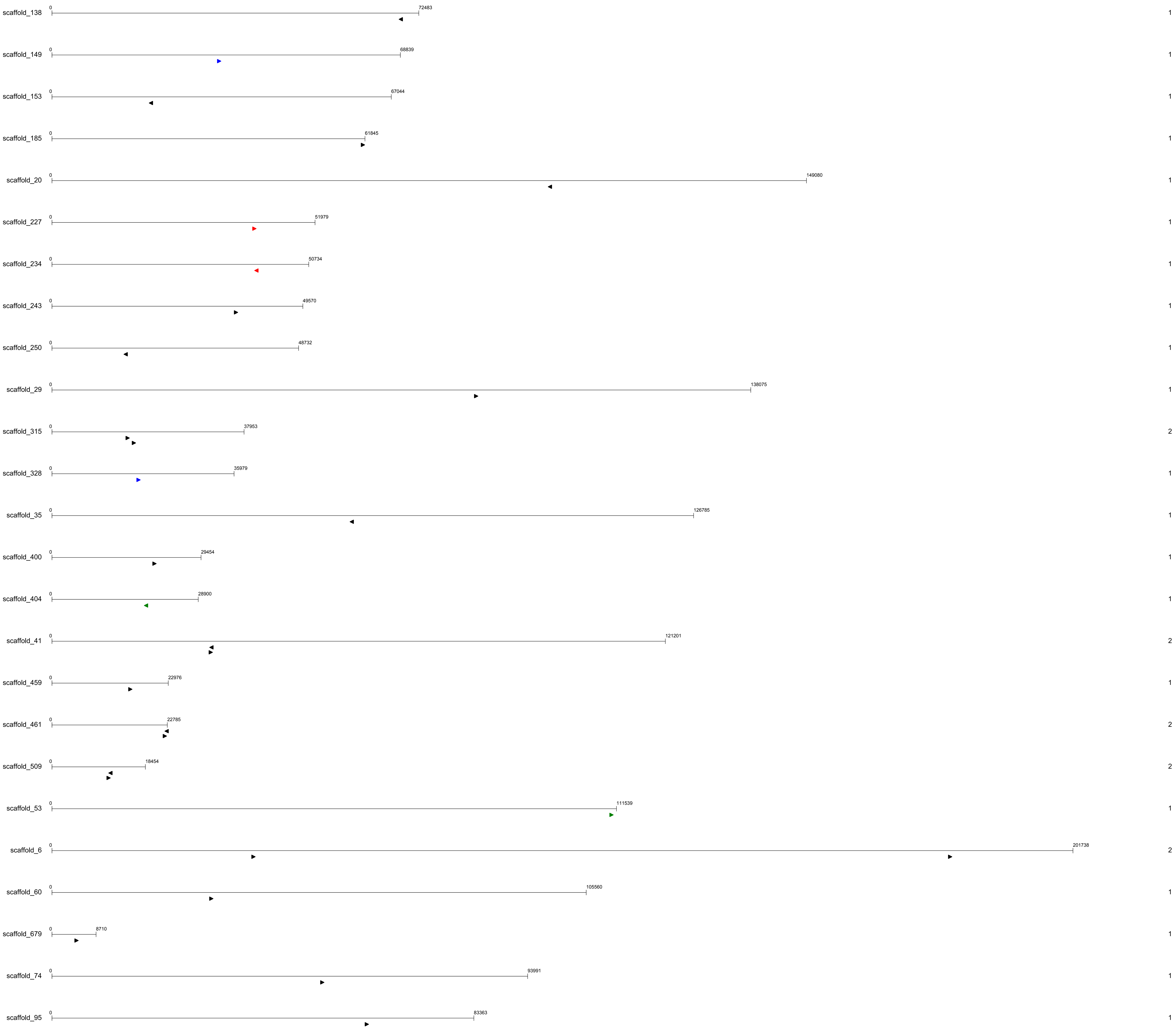

***P. violaceum* Class I RNAs (n=27)**

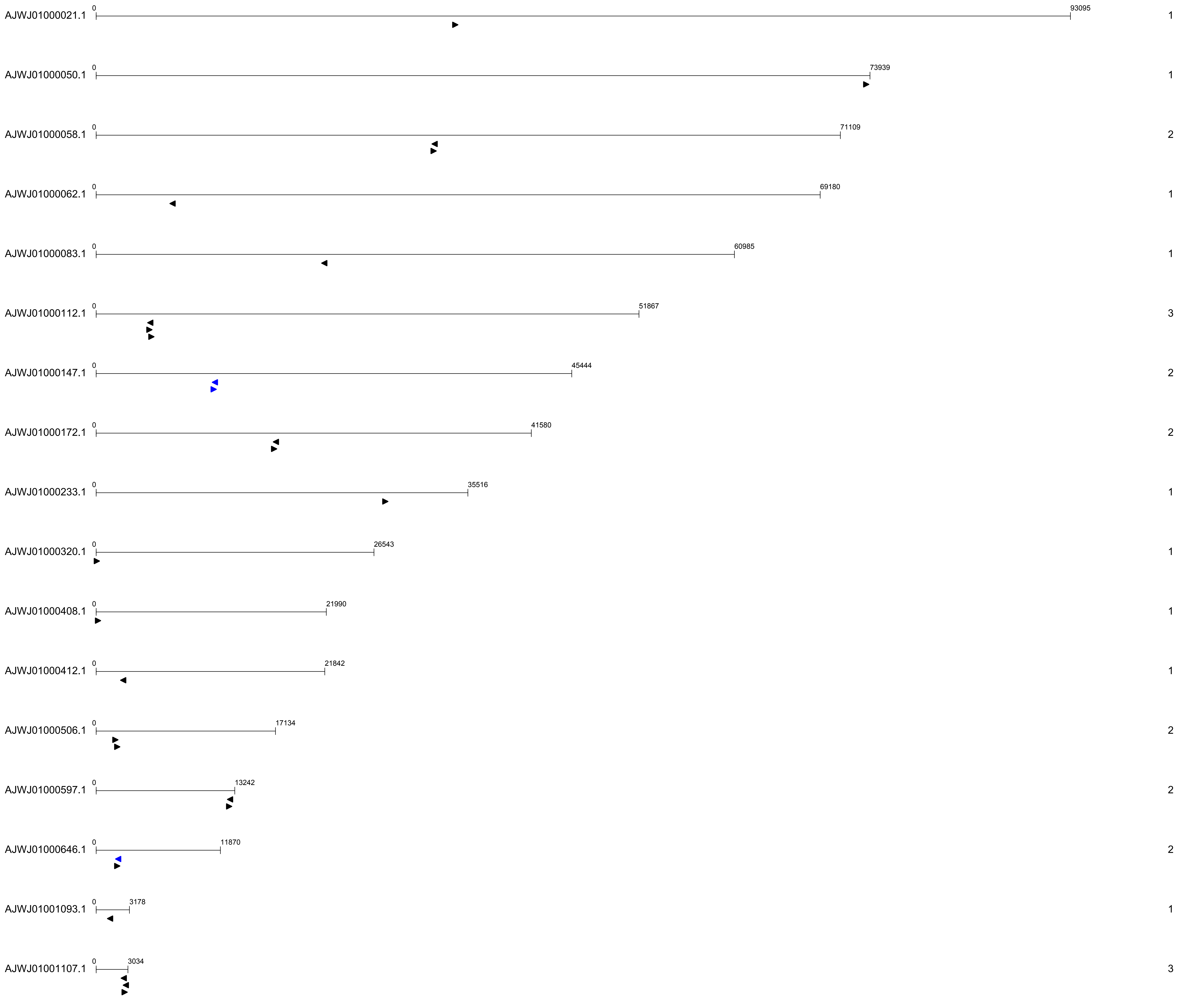

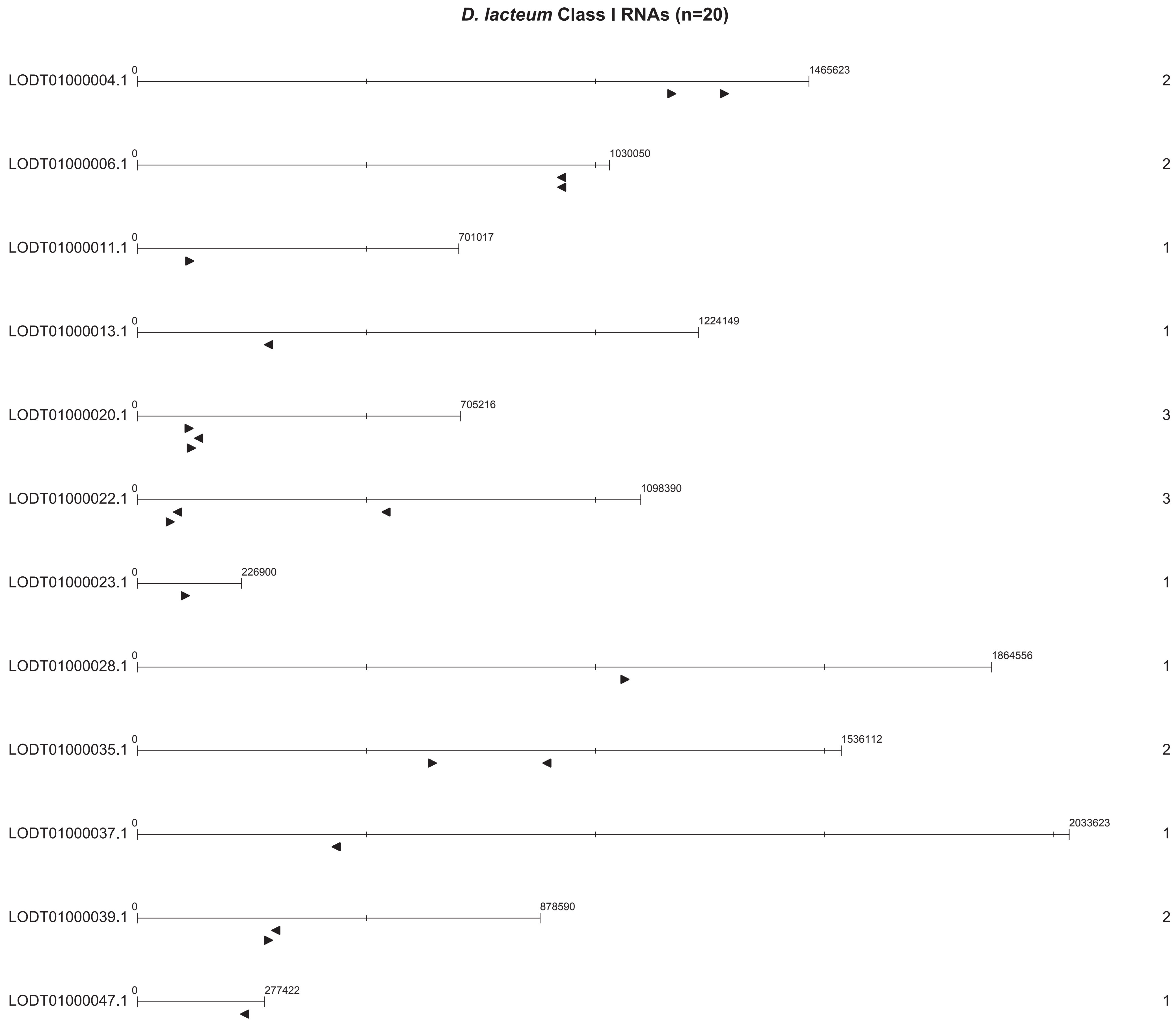
D. lacteum Class I RNAs (n=20)
LODT01000004.1
0
1465623
2
LODT01000006.1
0
1030050
2
LODT01000011.1
0
701017
1
LODT01000013.1
0
1224149
1
LODT01000020.1
0
705216
3
LODT01000022.1
0
1098390
3
LODT01000023.1
0
226900
1
LODT01000028.1
0
1864556
1
LODT01000035.1
0
1536112
2
LODT01000037.1
0
2033623
1
LODT01000039.1
0
878590
2
LODT01000047.1
0
277422
1

***D. caveatum* Class I RNAs (n=24)**

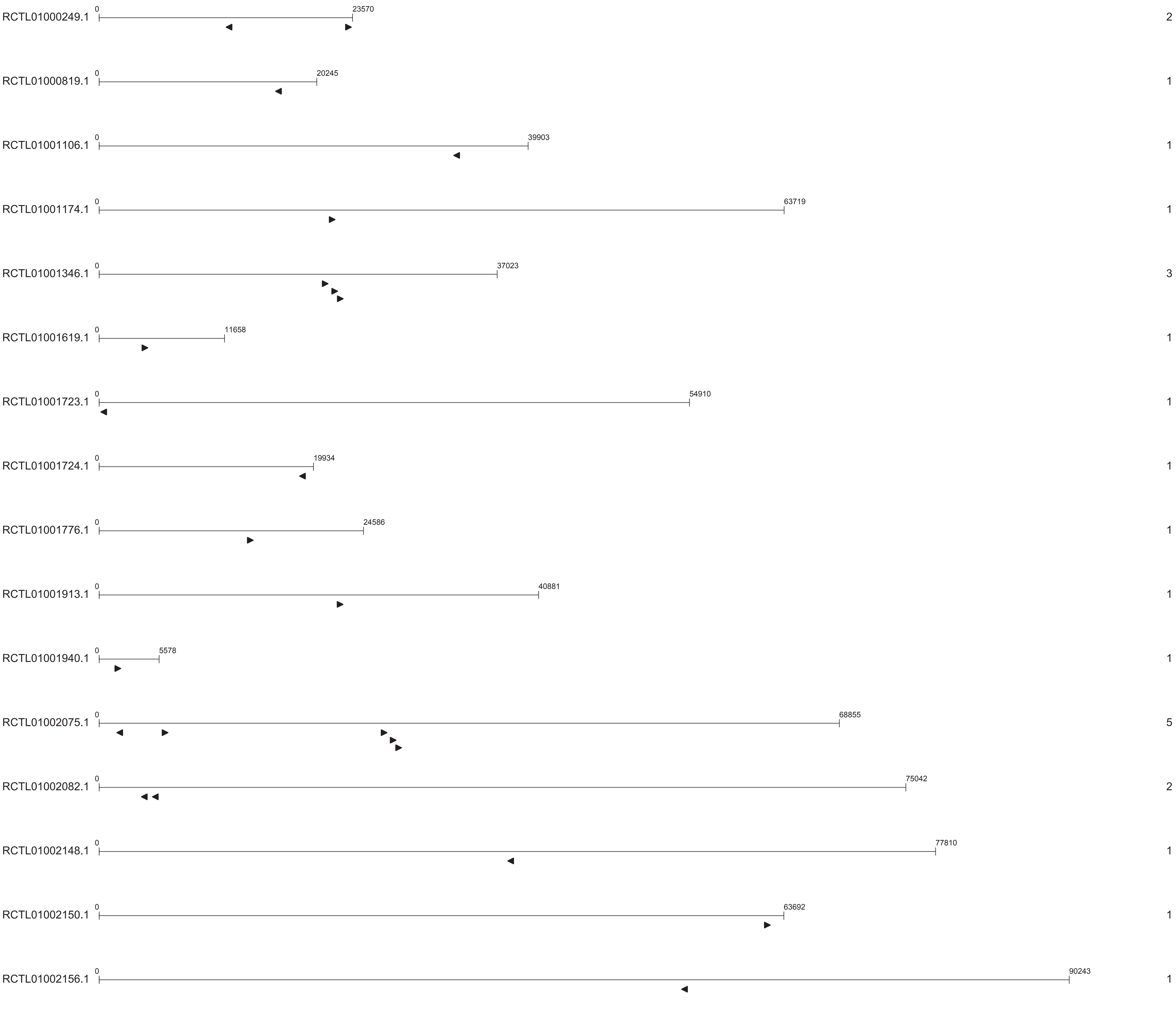

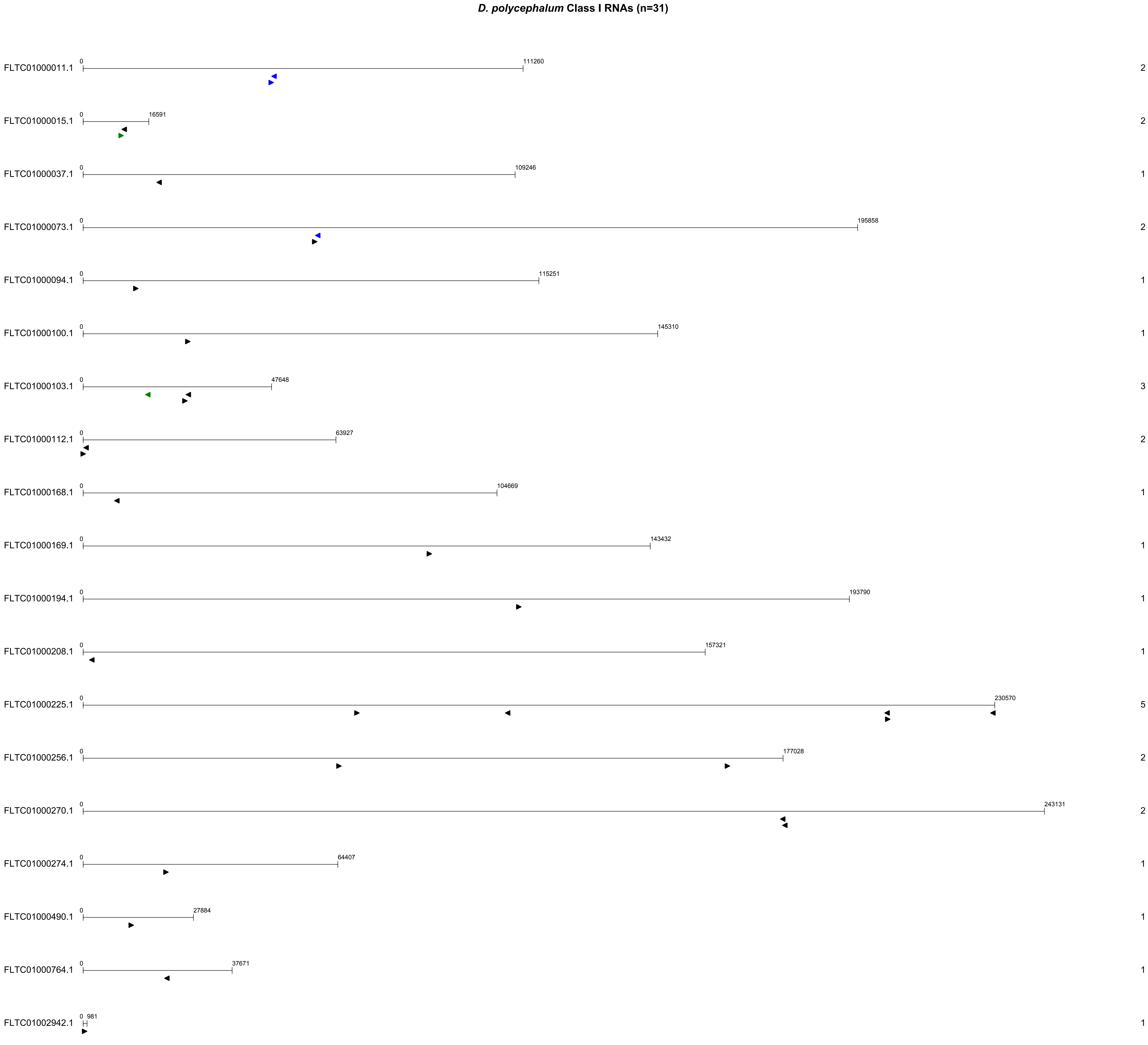

*D. polycephalum* Class I RNAs (n=31)
FLTC01000011.1
0
111260
2
FLTC01000015.1
0
16591
2
FLTC01000037.1
0
109246
1
FLTC01000073.1
0
195858
2
FLTC01000094.1
0
115251
1
FLTC01000100.1
0
145310
1
FLTC01000103.1
0
47648
3
FLTC01000112.1
0
63927
2
FLTC01000168.1
0
104669
1
FLTC01000169.1
0
143432
1
FLTC01000194.1
0
193790
1
FLTC01000208.1
0
157321
1
FLTC01000225.1
0
230570
5
FLTC01000256.1
0
177028
2
FLTC01000270.1
0
243131
2
FLTC01000274.1
0
64407
1
FLTC01000490.1
0
27884
1
FLTC01000764.1
0
37671
1
FLTC01002942.1
0
981
1

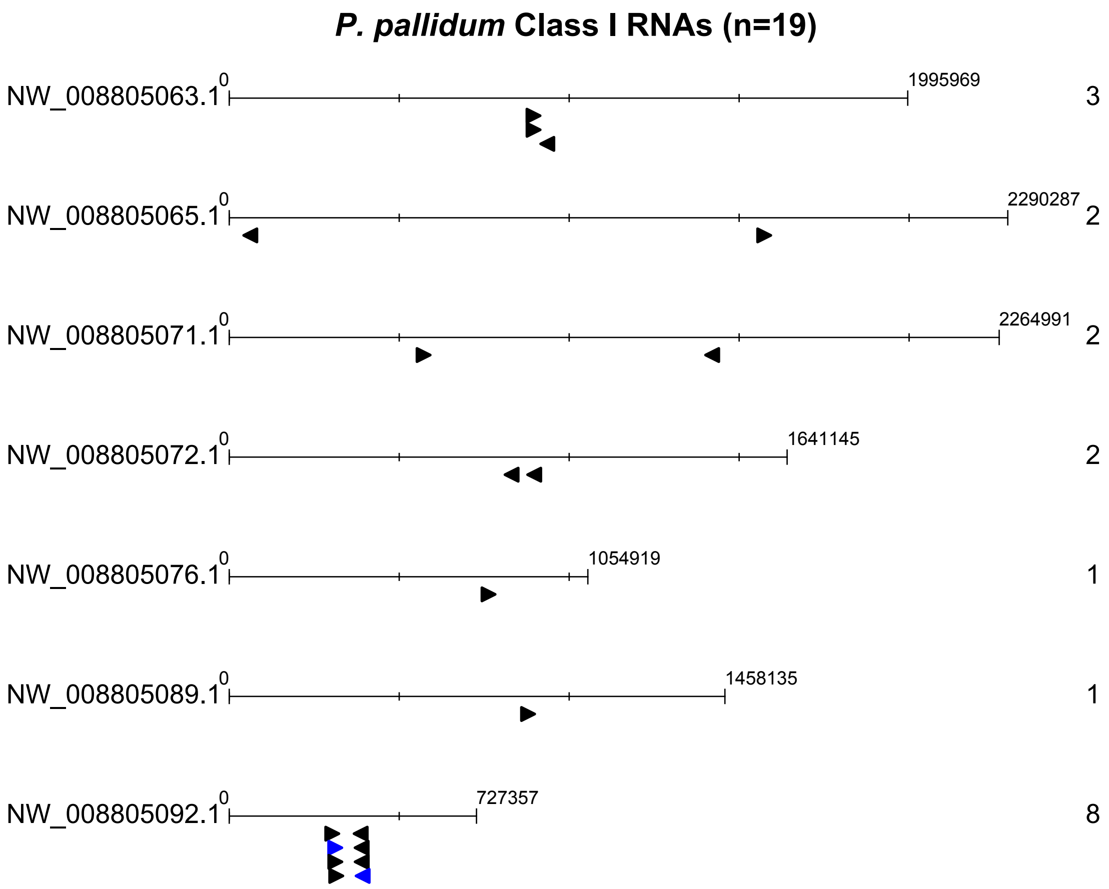

*P. pallidum* Class I RNAs (n=19)
NW_008805063.1
0
1995969
3
NW_008805065.1
0
2290287
2
NW_008805071.1
0
2264991
2
NW_008805072.1
0
1641145
2
NW_008805076.1
0
1054919
1
NW_008805089.1
0
1458135
1
NW_008805092.1
0
727357
8

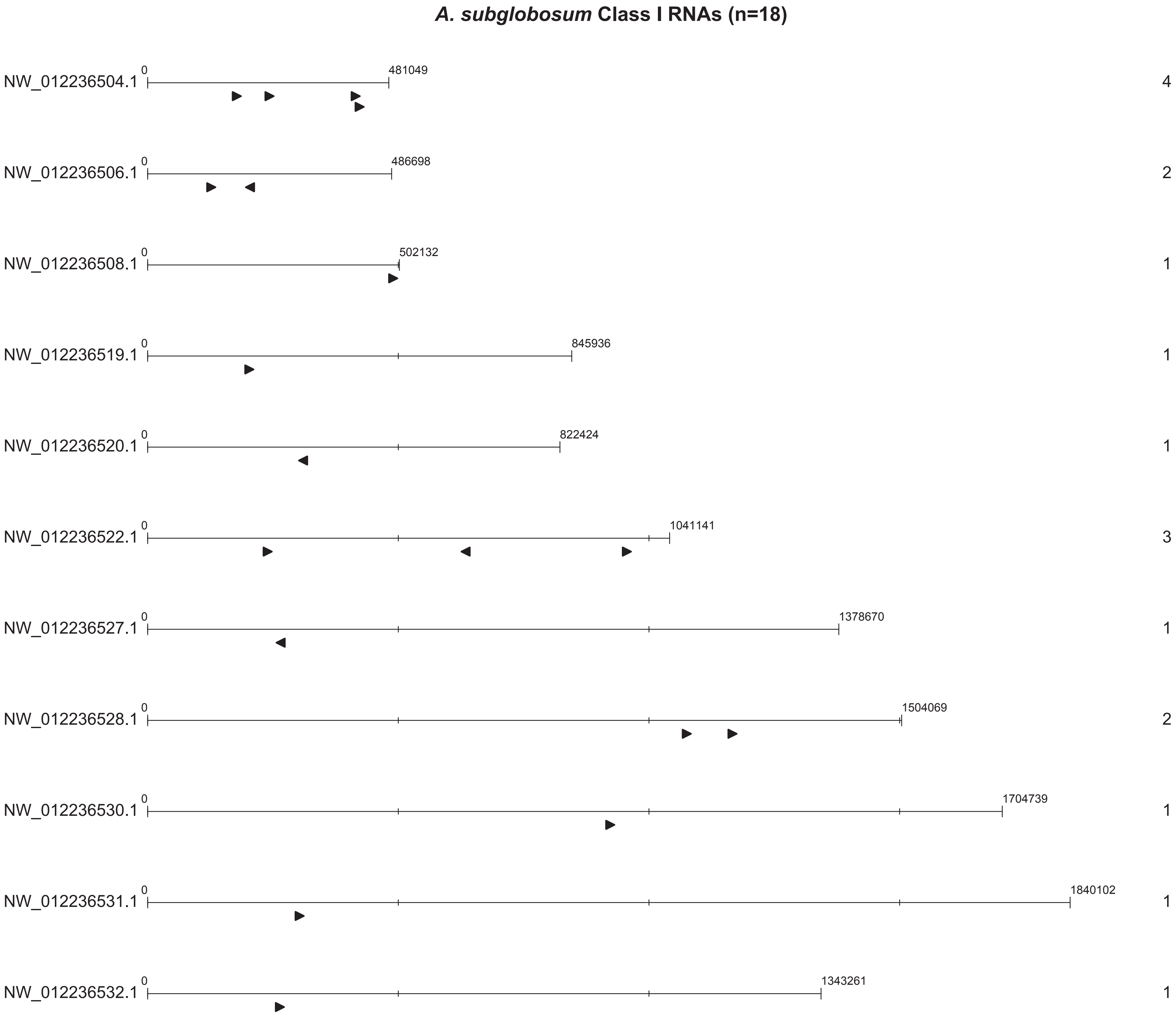

A. subglobosum Class I RNAs (n=18)
NW_012236504.1
0
481049
4
NW_012236506.1
0
486698
2
NW_012236508.1
0
502132
1
NW_012236519.1
0
845936
1
NW_012236520.1
0
822424
1
NW_012236522.1
0
1041141
3
NW_012236527.1
0
1378670
1
NW_012236528.1
0
1504069
2
NW_012236530.1
0
1704739
1
NW_012236531.1
0
1840102
1
NW_012236532.1
0
1343261
1

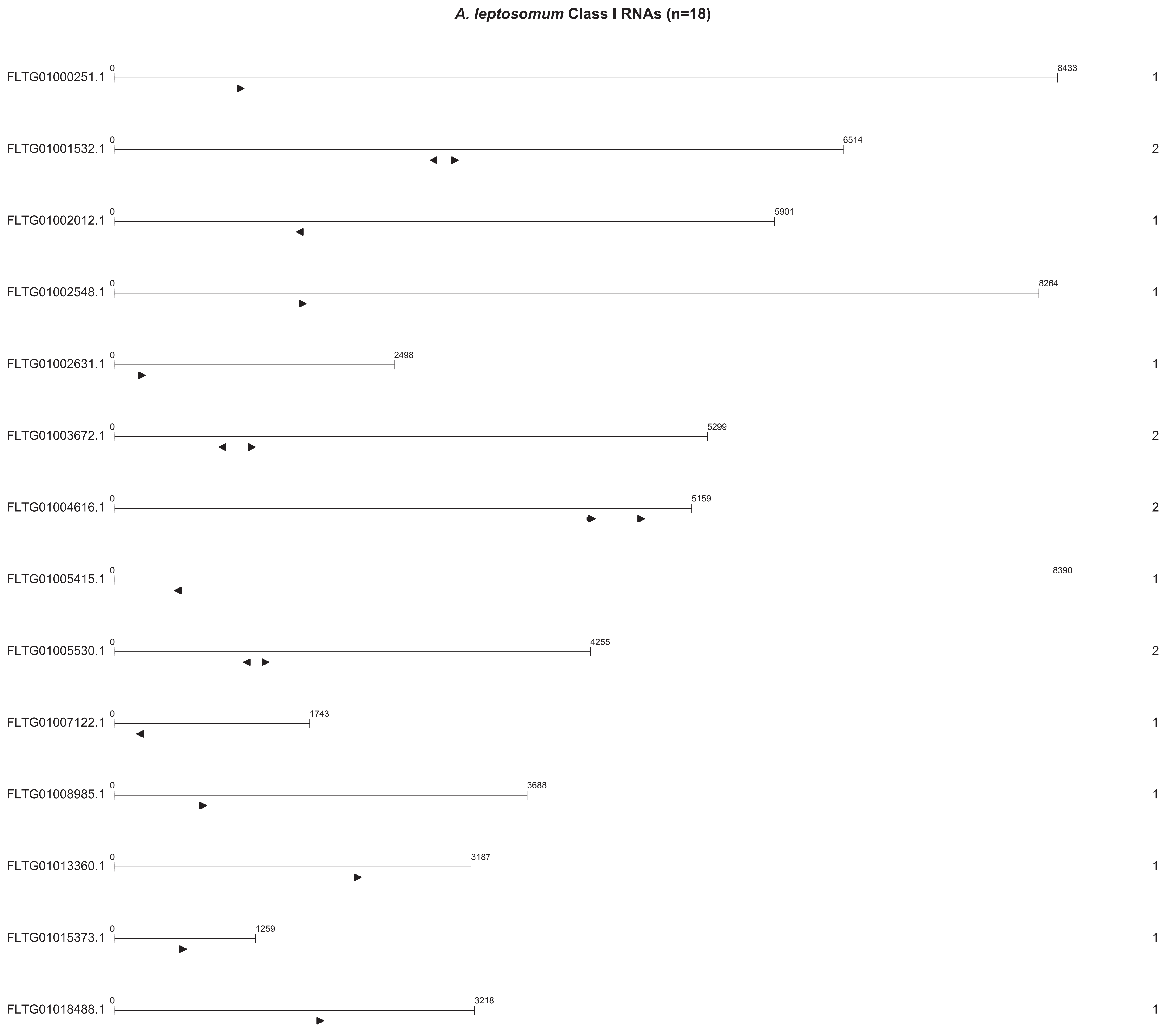
*A. leptosomum* Class I RNAs (n=18)
FLTG01000251.1 0 8433 1
FLTG01001532.1 0 6514 2
FLTG01002012.1 0 5901 1
FLTG01002548.1 0 8264 1
FLTG01002631.1 0 2498 1
FLTG01003672.1 0 5299 2
FLTG01004616.1 0 5159 2
FLTG01005415.1 0 8390 1
FLTG01005530.1 0 4255 2
FLTG01007122.1 0 1743 1
FLTG01008985.1 0 3688 1
FLTG01013360.1 0 3187 1
FLTG01015373.1 0 1259 1
FLTG01018488.1 0 3218 1

### *A. elipticum* Class I RNAs (n=17)

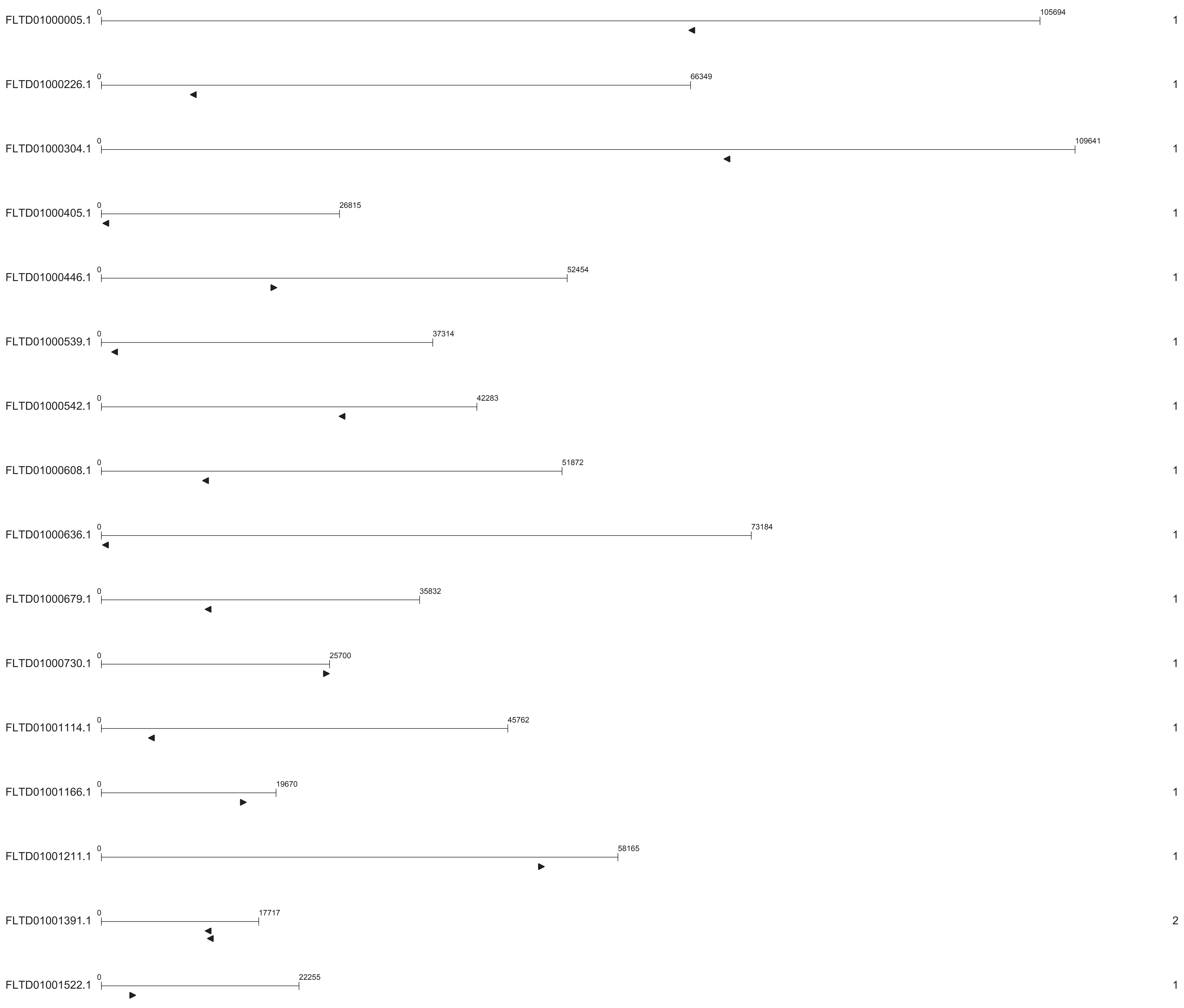

***D. polycarpum* Class I RNAs (n=25)**

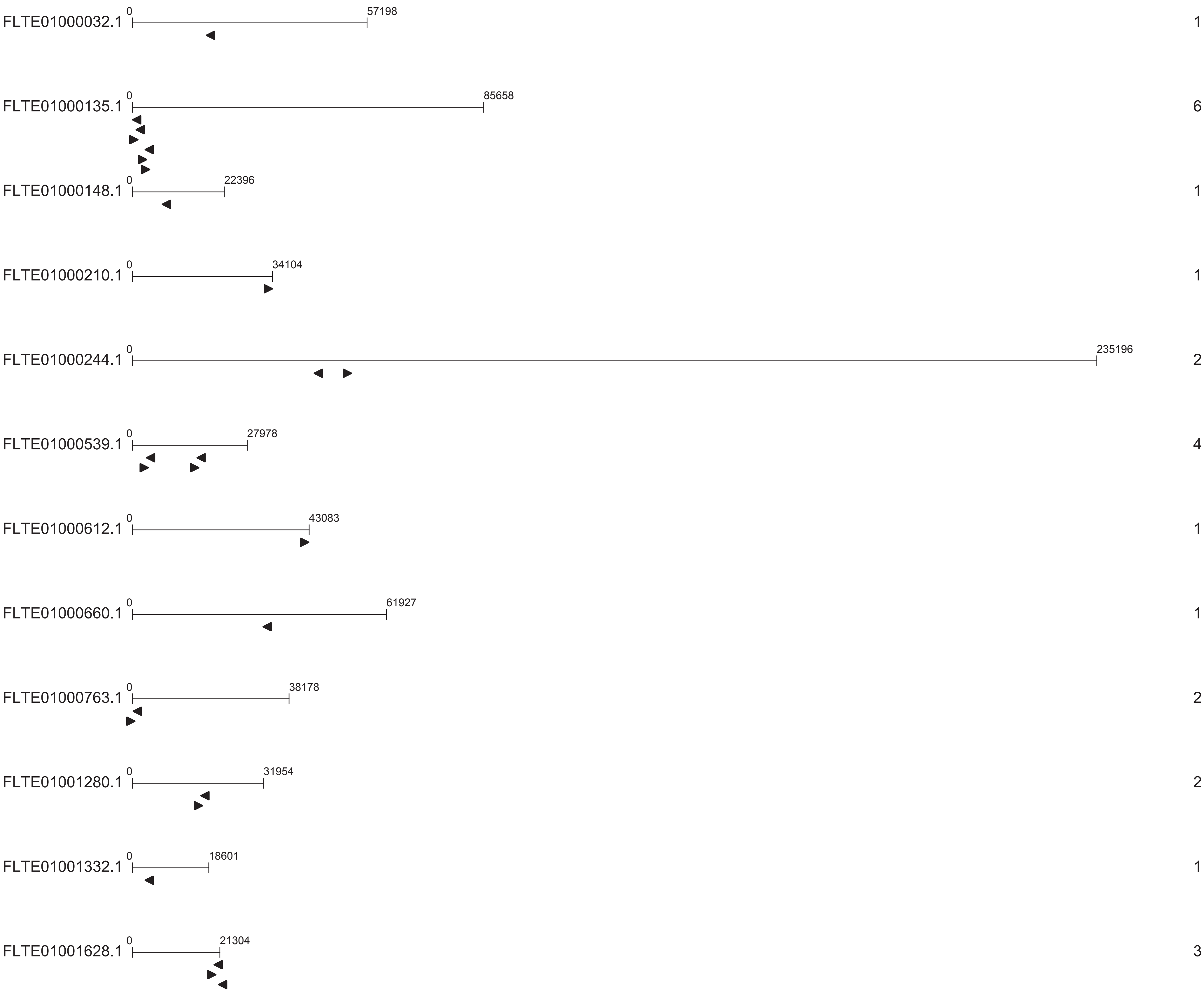

### *D. fasciculatum* Class I RNAs (n=19)

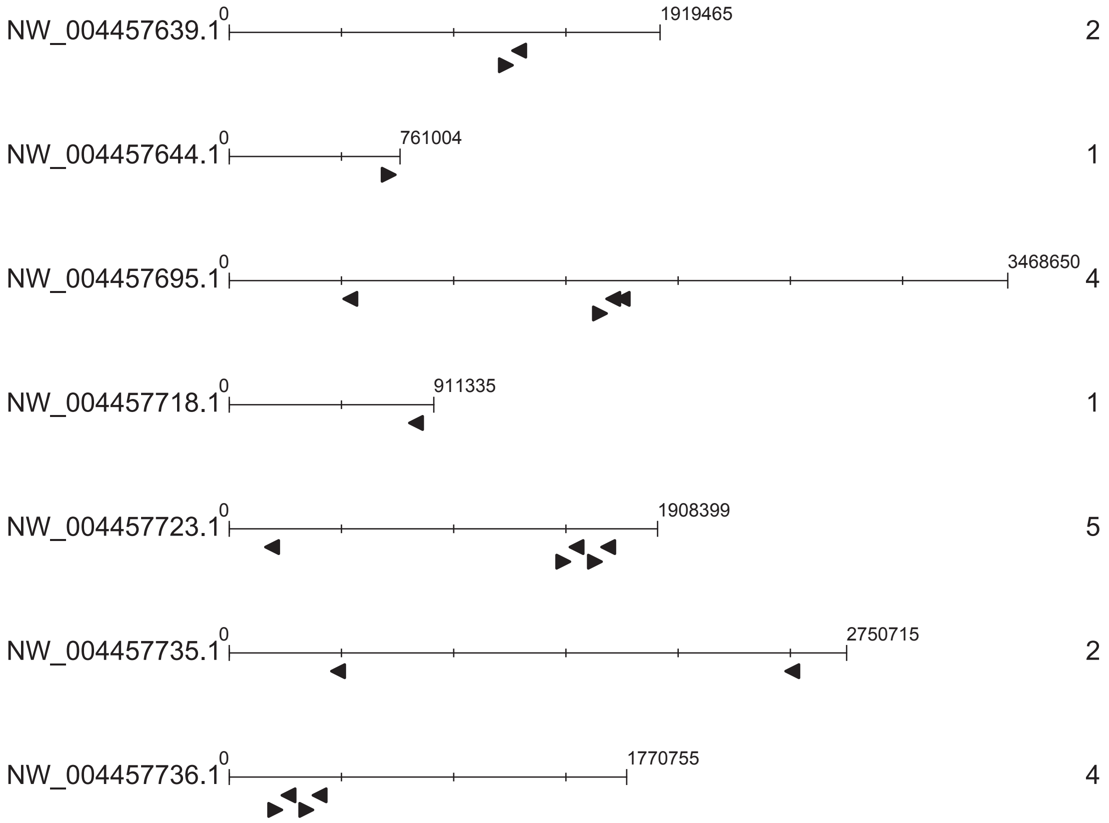

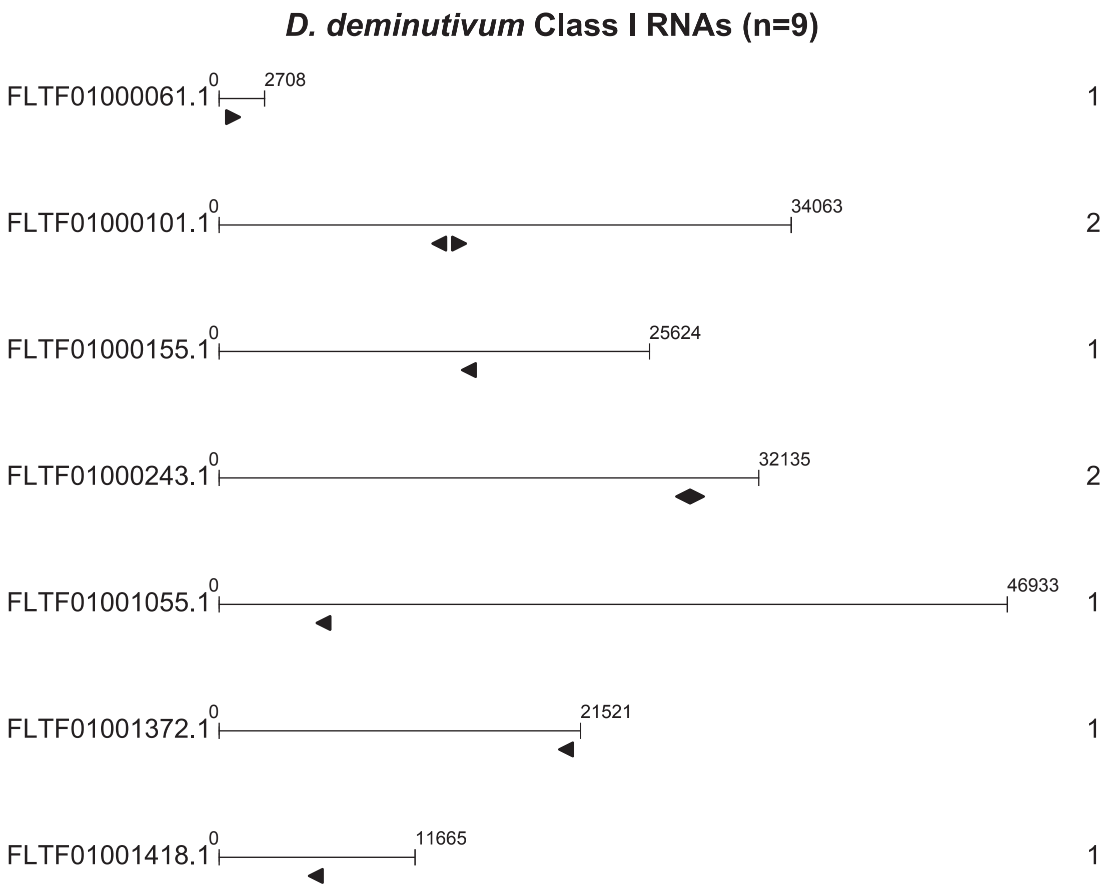
D. deminutivum Class I RNAs (n=9)
FLTF01000061.1
0
2708
1
FLTF01000101.1
0
34063
2
FLTF01000155.1
0
25624
1
FLTF01000243.1
0
32135
2
FLTF01001055.1
0
46933
1
FLTF01001372.1
0
21521
1
FLTF01001418.1
0
11665
1
