## Supplementary figures and images for "Abundantly expressed class of non-coding RNAs conserved through the multicellular evolution of dictyostelid social amoebae"

### Additional file 6

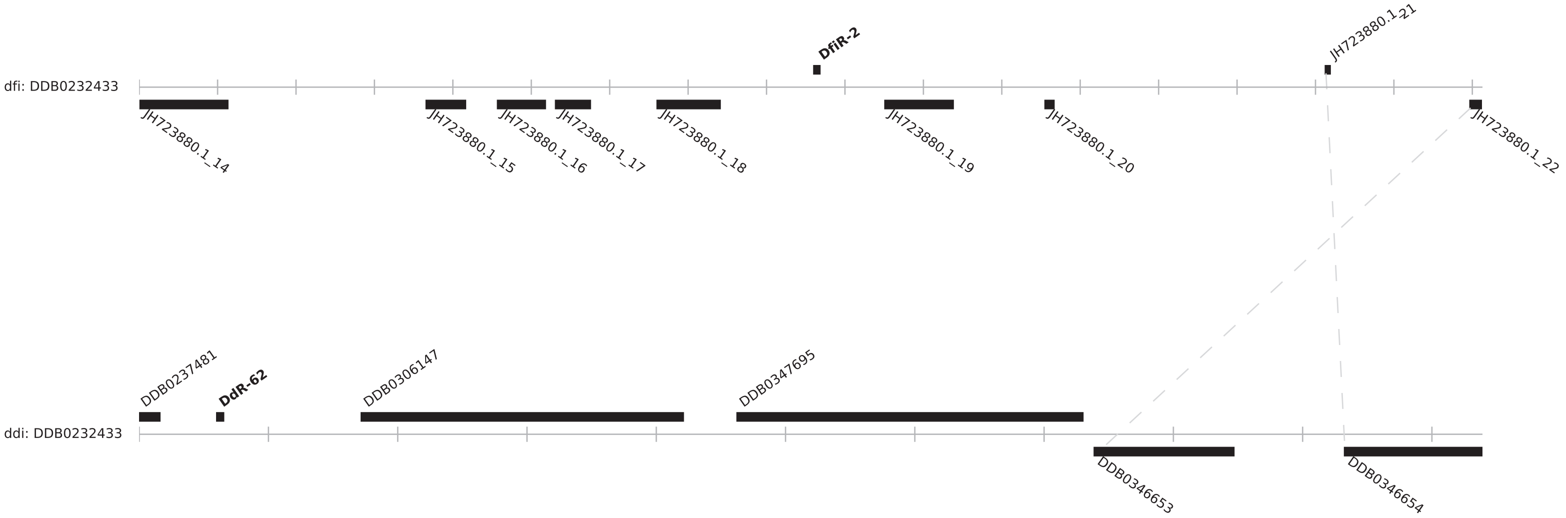

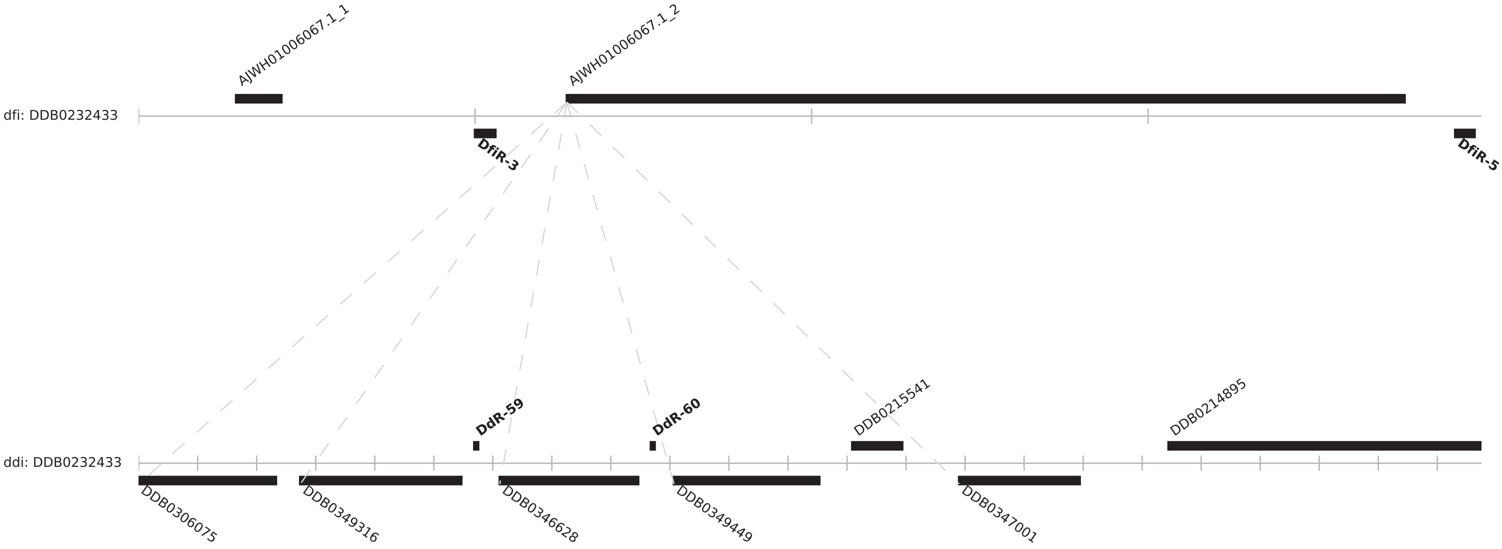

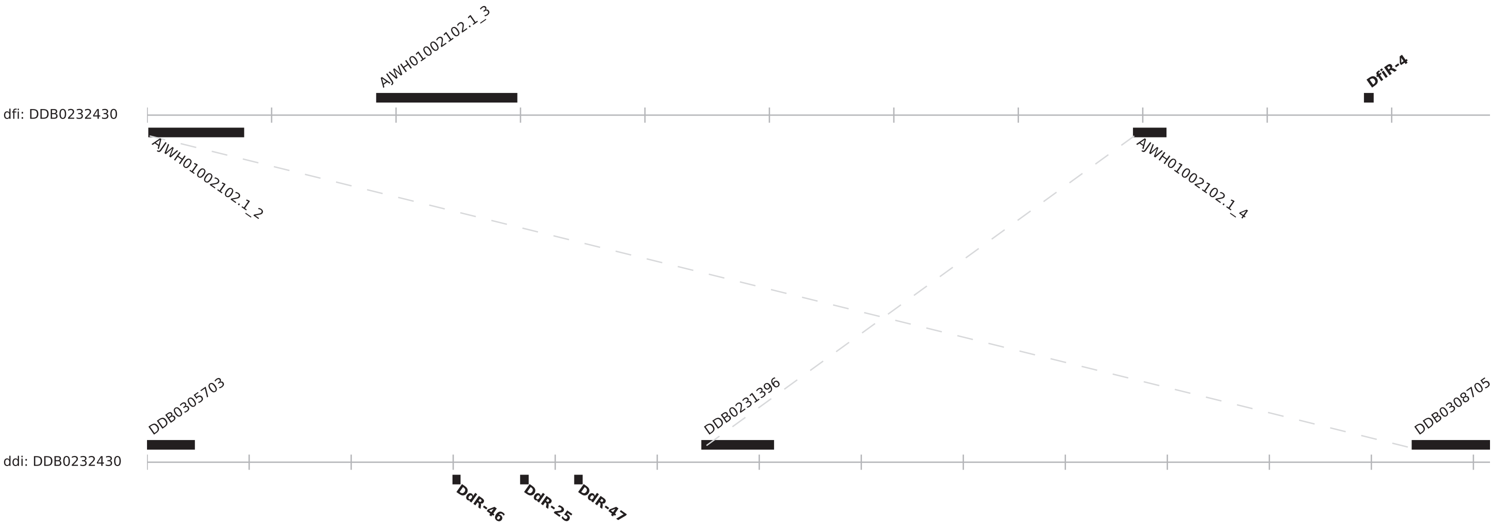

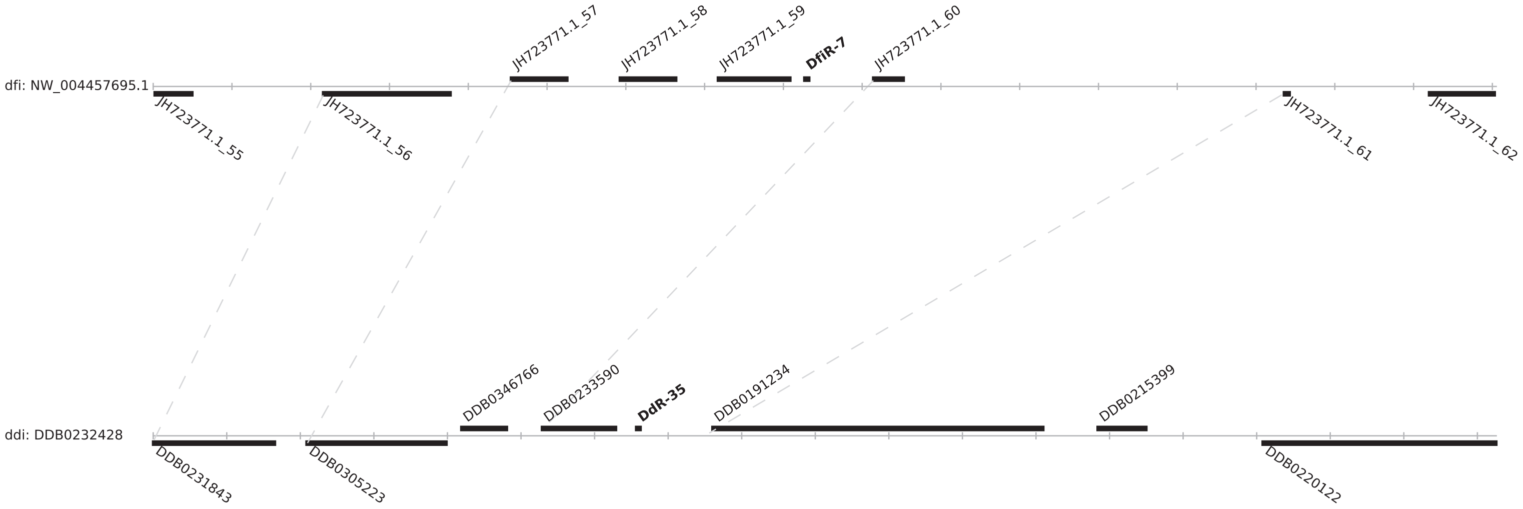

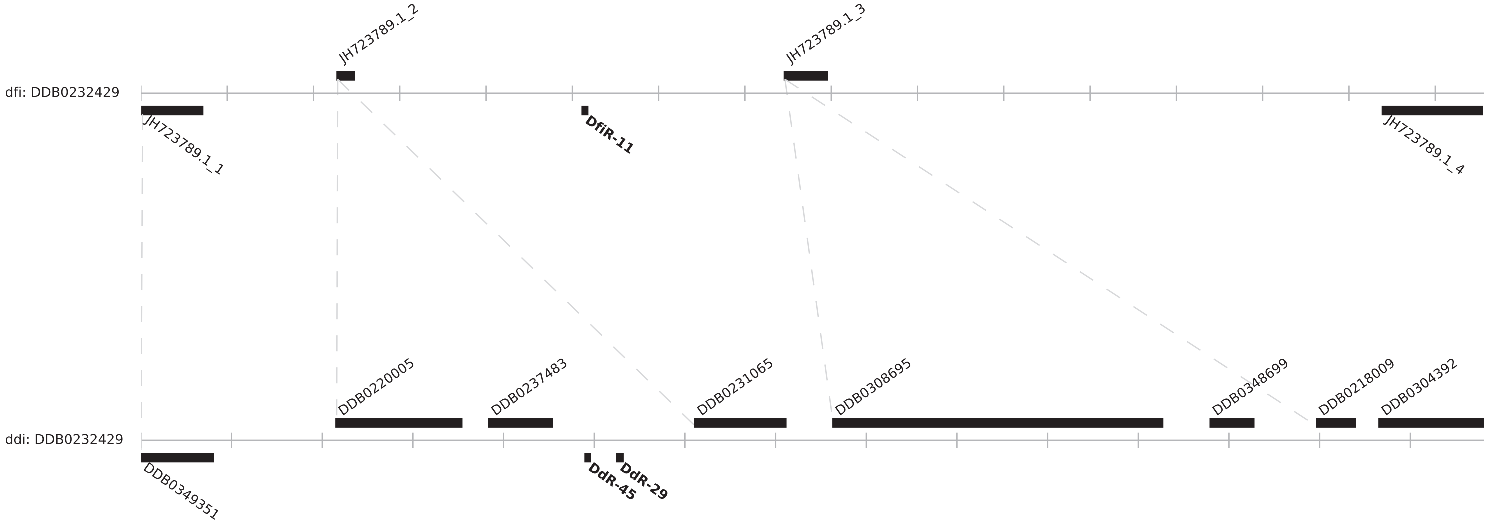
